# Rapid and consistent clustering of millions of genomes highlights the diversity of prokaryotic life

**DOI:** 10.64898/2025.12.30.695181

**Authors:** Johanna von Wachsmann, Leonie J. Lorenz, Matthew J. Russell, Tatiana A. Gurbich, Víctor Rodríguez-Bouza, Samuel T. Horsfield, John A. Lees, Robert D. Finn

**Author notes:** Contributed equally (John A. Lees, Robert D. Finn).

## Abstract

Bacterial genome and metagenome databases collectively contain over 5 million high-quality assemblies. However, the redundancy of these databases and the limited scalability of existing tools create bottlenecks for fully comprehensive, tree-of-life-scale genomic analyses. A fundamental task is to first break this data into smaller chunks, guided by their genome similarity. However, alignment-based comparative methods struggle to handle more than a few tens of thousands of genomes at a time, making the global organisation computationally complex and expensive. Here, we present *gemsparcl (*https://github.com/johannahelene/gemsparcl), a tool that clusters bacterial genomes into genomic cohesive units (GCUs), at approximately species-level resolution, over 500 times faster than existing methods. As part of developing *gemsparcl*, we developed sketchlib.rust, a one-permutation MinHash approach that implements an auxiliary inverted index to further accelerate all-versus-all comparisons. We added a statistical correction for incomplete metagenome-assembled genomes (MAGs) to enable accurate distance estimation and network-based edge quality filtering. After genome completeness quality control, we clustered 5.6 million high-quality bacterial genomes (2.88 million isolates and 2.77 million MAGs) into 92,954 GCUs in ∼14 hours using 48 CPU threads and less than 16.5 GB of memory. Using taxonomic validation of the GCUs, the method achieves very high (99.76%) cluster purity (meaning only one species label occurs per GCU). We demonstrate that the clustering also highlights cases where taxonomic naming can be potentially harmonised or improved. Furthermore, we identify the most frequently reconstructed MAGs that lack a corresponding isolate genome and are thus priorities for culturing. The enhanced speed of *gemsparcl* enables routine database updates to incorporate the latest genomes. It also makes reference-free microbiome analysis across millions of genomes computationally tractable for the first time.

## Introduction

Bacterial genomic databases continue to grow exponentially (Břinda *et al*. 2025), driven by two converging endeavours. First, the widespread adoption of whole genome sequencing in clinical and public health microbiology has made routine pathogen surveillance tractable, with organisms such as *Mycobacterium tuberculosis*, *Klebsiella pneumoniae*, and *Salmonella enterica* now sequenced at scale for outbreak investigation and antimicrobial resistance monitoring (Hunt *et al*. 2024; Timme *et al*. 2018). Second, advances in shotgun metagenomics have enabled the routine reconstruction of microbial genomes directly from environmental samples, bypassing the need for cultivation and unlocking enormous reservoirs of previously inaccessible diversity across a range of samples, such as soil, ocean, and host-associated microbiomes (Dmitrijeva *et al*. 2025; Gurbich *et al*. 2023; Schmidt *et al*. 2024). Together, these developments have resulted in the number of isolate genome collections quadrupling since 2021, with now millions of isolate genomes available (Hunt *et al*. 2024). Similarly, the number of metagenome-assembled genomes (MAGs) now runs into the millions. Current genomic databases contain massive collections: the AllTheBacteria (Hunt *et al*. 2024) database contains 2.7 million genomes, while combined resources from MGnify (Gurbich *et al*. 2023), SPIRE (Schmidt *et al*. 2024), and mOTUs (Dmitrijeva *et al*. 2025) contain over 7 million MAGs.

This huge volume of genomic data has created a fundamental organisational challenge for analytical scalability. Aggregating all genomes of a given species across surveillance projects, which is essential for robust epidemiological inference and antimicrobial resistance tracking, requires clustering at a scale that current tools cannot achieve. Also, identifying novel lineages within MAG datasets or tracking specific genomes in a One Health context demands methods that scale to millions of genomes without sacrificing biological resolution (Vallejo-Espín *et al*. 2025). Traditional bacterial taxonomy was largely established before genomic data became available, resulting in classifications that are sometimes inconsistent with evolutionary relationships (Parks *et al*. 2022). Alternatively, comparison of genome content can be used to infer the phylogenetic relatedness of a population without the need for taxonomic labelling (Lees *et al*. 2019). Clustering genomes by sequence similarity therefore captures biologically meaningful evolutionary distances, independent of the historical contingencies embedded in existing nomenclature. Genomes clustered by sequence similarity (or GCUs, genomically cohesive units) represent a principled basis for pangenome analysis.

After clustering, a common next step in epidemiological or evolutionary analysis is dereplication, in which a single representative genome is chosen for each cluster, typically based on genome quality metrics or genome similarity (e.g. the cluster centroid). These representative genomes then serve as delegates for the other genomes, resulting in a dataset reduction used in downstream analyses. These non-redundant genomes can then be used to accelerate searches, avoiding the need to search the entire collection. Clustering also lays the foundation for addressing broader questions about species delineation and microbial diversity.

However, with genomes now numbering in the millions, current tools cannot keep pace with the computational challenge of genome clustering, which centres on determining evolutionary relatedness between pairs of bacterial genomes within a collection. Two genomes sharing an average nucleotide identity (ANI) ≥ 95% are typically classified as belonging to the same species (Caro-Quintero and Konstantinidis 2012). Traditional clustering approaches, such as dRep (Olm *et al*. 2017), which uses genome-wide ANI (gANI; (Varghese *et al*. 2015) for distance estimation, rely on whole-genome sequence alignment, resulting in computational complexity that scales quadratically with dataset size. While manageable for datasets containing thousands of genomes, these methods become prohibitively expensive for current collections that far exceed that number. Clustering millions of genomes using such approaches would result in processing times that extend to months and memory requirements that surpass typical available computational resources (Almeida *et al*. 2021). Alternatively, sketching algorithms have emerged as a solution to address this scalability issue (Rowe 2019). Rather than performing costly whole-genome alignments, sketching methods create compact genomic “fingerprints” by selecting a small, representative subset of genomic k-mers (short DNA sequences of length k) and converting them into numerical hashes. Tools such as mash (Ondov *et al*. 2016) pioneered this approach, and sourmash (Pierce *et al*. 2019) extended it, enabling rapid pairwise comparisons through MinHash (Broder *et al*. 2000) sketching techniques that approximate ANI values with high accuracy near the species boundary, at around 95%. This approach preserves essential similarity information while requiring orders of magnitude fewer computational resources than full-sequence alignments, since typically only 1000 k-mers are sufficient (compared to >1M in a typical bacterial genome).

The integration of MAGs into large-scale genomic analyses introduces additional complexity that current methods do not address. While MAGs are a crucial resource for understanding unculturable bacterial diversity (Mirete *et al*. 2025), these genome assemblies are often incomplete and/or contaminated. This incompleteness creates a systematic bias: when MAGs are compared to complete genomes from the same species, the reduced genomic overlap (including k-mer-based approaches) artificially decreases similarity scores, thereby underestimating true relatedness and potentially fragmenting species clusters (see Supp. Figure 2).

Despite ongoing method development efforts across the research community, no existing tool addresses the combined challenges of scale, speed, and MAG-specific artefacts. Established dereplication tools, such as dRep, become computationally challenging at tens of thousands of genomes, necessitating laborious dataset partitioning and hierarchical clustering approaches that still scale poorly to larger datasets (Almeida *et al*. 2021). Here, we present *gemsparcl*, a novel sketching-based clustering approach that addresses these fundamental limitations through three key innovations. First, binned sketching with an inverted index restricts distance calculations to genomically similar sequences, achieving sub-linear runtime scaling. Second, explicit correction for genome incompleteness enables accurate distance estimation for incomplete genomes alongside complete genomes. Third, k-nearest neighbour filtering focuses edge creation on biologically relevant comparisons, enabling efficient sparse network construction at database scale.

## Methods

### Sketching genomes

The sketching process used in *gemsparcl* is the same as that previously described in Mandrake (Lees *et al*. 2022), and illustrated in Figure 1. Briefly, the sketching process begins by extracting k-mers from the input genome sequences and applying ntHash2 (Kazemi et al. 2022) to convert them into 64-bit numerical values. We employ one-permutation MinHash (Li *et al*. 2012), which applies a single permutation to the k-mer feature space of the genome and divides it into s buckets of equal size (where s equals the desired sketch size). For each bucket, we identify the minimum hash value among all k-mers falling within that bucket’s range. To minimise memory usage, we store only the 14 most significant bits of each minimum hash value, rather than the full hash, which substantially reduces the sketch size. This is the same approach as BinDash (Zhao 2019), but we reimplemented the algorithm in Rust and confirmed that the sketches were identical to those in the C++ implementation.

**Figure 1:**
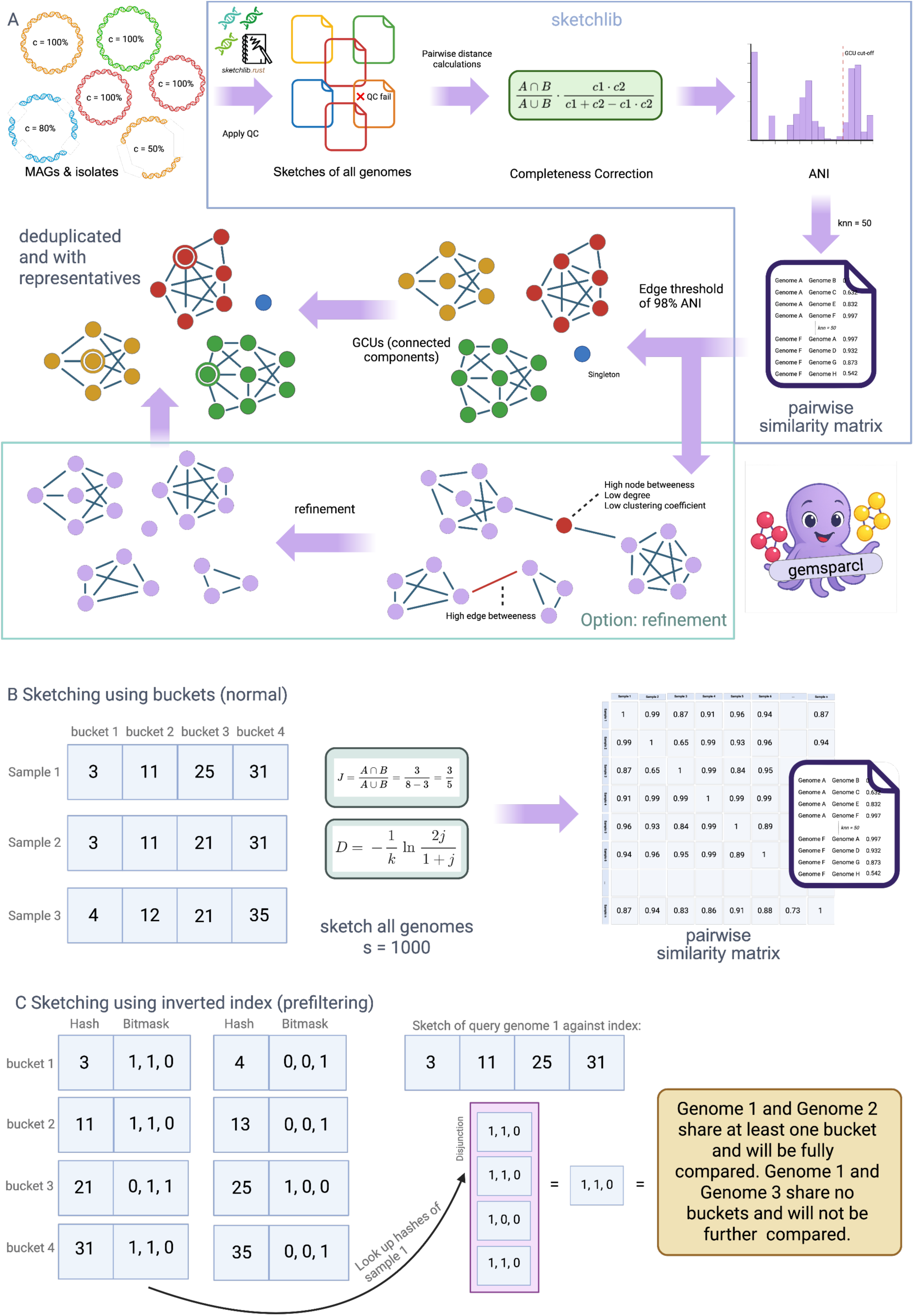
Overview of the main *gemsparcl* method: (A) Bacterial genomes are taken as input, including both isolate genomes sequenced from pure cultures and metagenome-assembled genomes (MAGs) reconstructed from environmental sequencing data. All genomes must pass defined quality thresholds (≥80% complete, <5% contaminated) before proceeding. The remaining genomes are handed to sketchlib.rust for sketching. Rather than comparing full genome sequences directly, which would be impossibly slow at scale, each genome is first compressed into a compact numerical fingerprint called a sketch. This is done by extracting all short-sequence substrings of length k (k-mers), hashing each to a numerical value using nt-Hash2, and then applying one-permutation MinHash. The full range of possible hash values is split into *s* equally sized buckets, and only the smallest hash value in each bucket is retained. The result is a fixed-length array of *s* numbers that captures the sequence composition of a genome in a fraction of its original size. Pairwise genomic distances between all genomes are then estimated from these sketches as approximate average nucleotide identity (ANI) values. For MAGs, an additional correction step is applied: incomplete genomes share proportionally fewer k-mers than complete ones due to missing sequence, which would otherwise cause them to appear more distantly related than they actually are. Correcting for completeness brings their estimated distances in line with those of complete genomes (where *c*_1_ and *c*_2_ are the estimated checkM/checkM2 completeness scores). The result is a pairwise ANI matrix that captures the genomic relationships among all input genomes. This matrix is used to build a similarity network, where each genome is a node and an edge is drawn between two genomes if their ANI is at least 98%. Connected groups of nodes within this network define genomically cohesive units (GCUs). These are sets of genomes that are all linked to one another through a chain of highly similar connections. GCUs are clusters of genomically similar sequences and need not follow existing taxonomic boundaries. From these clusters, a representative genome (currently, 1 of 500 is selected) can be selected per GCU (prioritising the most complete genome), and redundant sequences, including exact duplicates with 100% identity, can be removed. Optionally, a refinement step can be applied to remove nodes or edges that act as weak bridges between otherwise distinct clusters. For example, contaminated genomes whose sequences span two lineages can be removed using the --refine flag. (B) Sketching algorithm. The sketching approach is based on BinDash. All k-mers from a genome are hashed in a single pass, and the full space of hash values is partitioned into a fixed number of buckets. The smallest hash value in each bucket is kept as that bucket’s representative, giving a compact, fixed-size sketch per genome. Because each k-mer is hashed only once and binning is performed immediately, this step is very fast, even for large datasets. Jaccard similarity is then calculated between pairs of sketches, counting how many buckets share the same minimum hash, and converted to an ANI estimate. The resulting pairwise values are written to a distance matrix. (C) Inverted index prefiltering. For very large datasets (e.g. >100k genomes), computing all pairwise distances, even from sketches, can be expensive. An inverted index is used as a prefiltering step to avoid wasting time on genome comparisons that are clearly not similar. First, all genomes are quickly sketched at low resolution (s=10 buckets). An inverted index is then built from these small sketches: for each bucket position, the index records which genomes share the same minimum hash in that bucket. Any pair of genomes that does not share at least one bucket in this index is extremely unlikely to be similar and is skipped entirely. Full-resolution sketches (s=1000) and precise distance calculations are performed only for genome pairs that pass this filter, thereby reducing total computation without affecting the final results.

To store sketches, we use a simple flat-file structure, writing the bits of each sketch bucket serially. This allows random access to sketches indexed by bucket index, k-mer index and genome index. Sketch metadata (e.g. sample name, data quality) is stored in a separate serialised file. This design has the following advantages: (1) the sketch file can be compressed efficiently using standard tools (snap (Zaharia *et al*. 2011), by default); (2) the file can be memory mapped, so when a subset of sketch data is being processed only the samples used are loaded into RAM; (3) merging sketch files simply appends data; (4) a single block of memory is used, avoiding pointer chasing; and (5) low-RAM parallelisation is implemented by having a master thread writing to file while worker threads sketch individual samples.

### Computing between-genome distances

Following sketch generation, pairwise distances between genomes are computed from the resulting hashes by counting the number of buckets that share the same minimum hash for pairs of samples. This gives the Jaccard similarity, where the matching bucket represents *S*1 ∩ *S*2, and since both sketches have the same size *s*, the denominator simplifies to 2*s* - bucket matching:

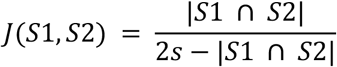

Distance metrics can be reported as either Jaccard distances (ranging from 1 {no shared k-mers} to 0 {identical genomes}) or as estimated ANI value calculated from the underlying Jaccard similarities J using the Mash distance:

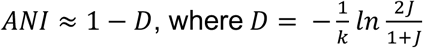

This is parallelised over all comparisons of the upper triangle of the pairwise distance matrix.

For large numbers of genomes, some distances must be discarded as there are too many to retain in RAM and writing to disk would be prohibitively slow. Therefore, only the 50 nearest neighbours (those with the highest Jaccard similarity) are kept in a priority queue. This is done on a per-sample basis and disregards exact matches, so the top 50 distances from sample A do not necessarily reciprocally include all samples which have sample A in their top 50. Although this breaks reciprocity, it allows for much more efficient computation and memory use.

### Inverted index for accelerated clustering

To enable efficient all-versus-all comparisons for large datasets, we implemented an inverted index data structure over genome sketches. This approach transforms pairwise comparison from O(N²) complexity to output-sensitive complexity, where query time depends on the number of similar genomes rather than total database size. This approach avoids computing full distances between genomically dissimilar sequences, reducing the number of comparisons from all possible pairs to only those likely to exceed the similarity threshold, thereby achieving an order-of-magnitude speedup over traditional methods and reaching sub-linear runtime (Supp. Figure 3). Recent theoretical work has shown that inverted indexes of sketch fingerprints can achieve such output-sensitive searches while maintaining the same space complexity as standard forward sketch representations (Ingels *et al*. 2025).

The inverted index uses the binned structure of our sketching algorithm to create a lookup table that maps hash values to the genomes that contain them. For each sketch bucket position (0 to s-1), we maintain a hash table with k-mer hashes as keys, and values of 0 and 1 to indicate which genomes have this minimum hash. PhyAlign (Břinda et al. 2025) demonstrated that ordering samples in a bitvector such that genetically similar samples are adjacent substantially improves compressibility. We implement this by allowing sorting based on external labels (using species labels) and by storing bitvectors as RoaringBitmaps (Lemire *et al*. 2017). In Supplementary Figure 1, we show the reduction in file size due to reordering.

Construction requires a single pass through all genome sketches, populating the index in O(N×S) time where N is the number of genomes and S is the sketch size. For the largest datasets, we employ a two-stage approach analogous to seed-based prefiltering methods like MMseqs2 (Steinegger and Söding 2017). Initial screening uses an inverted index of extremely small sketches (s = 10 buckets) to rapidly identify candidate genome pairs sharing any k-mer content. This dimensionality reduction, using sketches 100× smaller for initial screening. However, unlike the amino acid comparisons performed by MMseqs2, where almost all pairwise comparisons lack shared sequence content and can be excluded, a relatively large proportion of genome comparisons (∼5%) share content due to database bias and conserved sequences.

We tested a random subset of 10k genomes from AllTheBacteria dataset with k=21, 61, 101, 301 and s=3, 5, 8, 10, 20, ensuring either at least one, all but one, or all buckets must match to pass the prefilter (Supp. Table 1). Using s=10 and k=21, with at least one bucket matching, was the smallest sketch size that gave no false negatives (i.e., sample pairs filtered out incorrectly) and removed 87% of distance comparisons (the best case would be removing 97.5% of comparisons). The two-stage approach therefore reduces the number of full sketch comparisons from O(N²) to O(M) (M << N), where M denotes the number of pairs sharing k-mer content, which is typically orders of magnitude smaller than N². Based on this, for each genome, we first query the sparse index in O(Ssmall) time to identify genomes that share at least one hash value across the 10 bucket positions. Only genome pairs passing this prefilter undergo full distance calculation using standard-sized sketches (s = 1000 buckets).

**Table 1:**
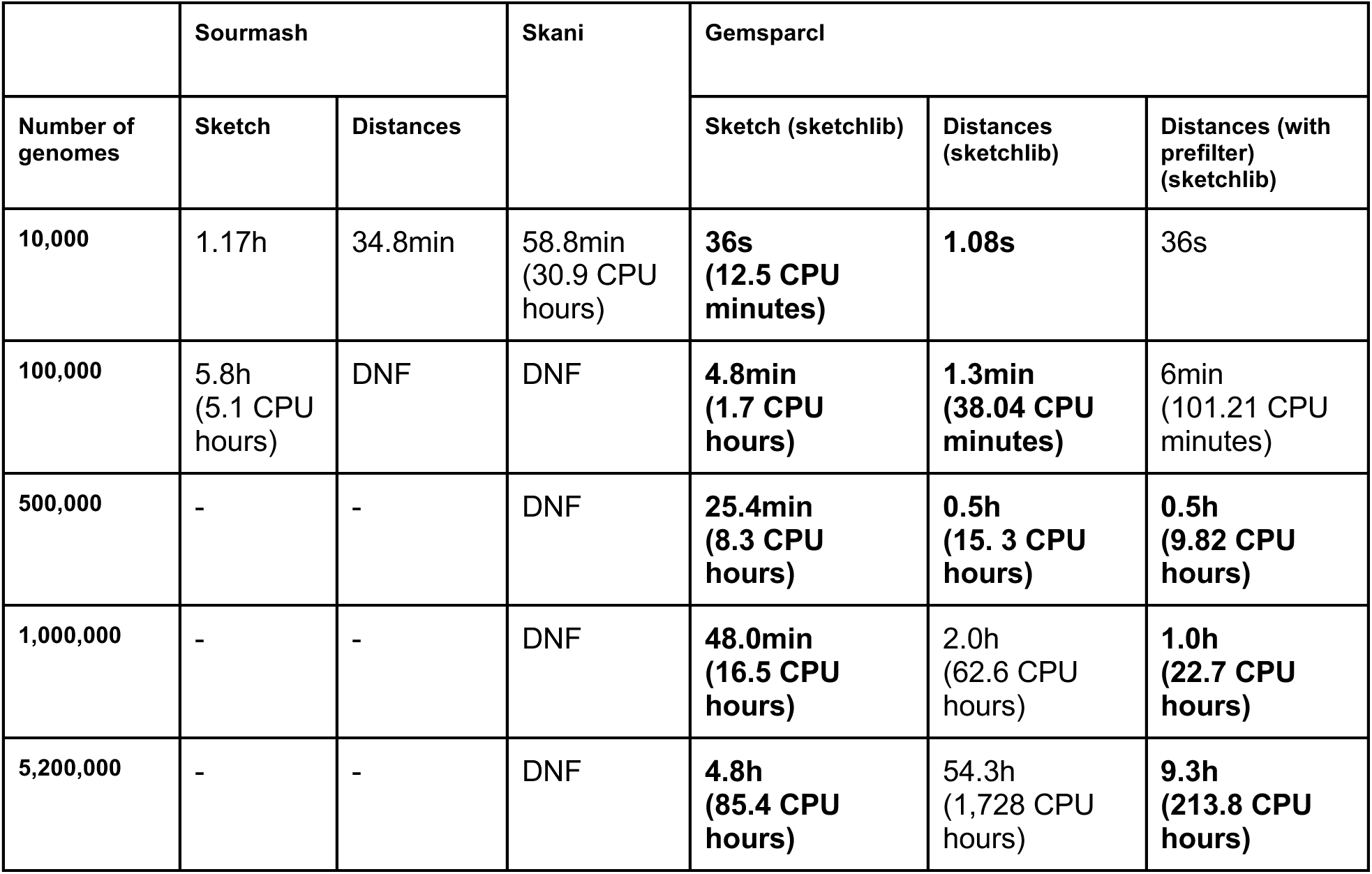
Wall-clock runtime of pairwise genome similarity tools across dataset scales. Benchmarking was performed on an Intel Xeon Gold 6336Y processor (2.40–3.60 GHz, 32 threads). CPU hours are shown in parentheses. Sketchlib separates the workflow into a sketching step and a distance-calculation step, with the latter optionally using a prefilter to accelerate large-scale comparisons. DNF indicates the job was killed after 48 hours. Dashes indicate tools that could not be evaluated at that scale due to prior failure on a smaller dataset.

|  | Sourmash |  | Skani | Gemsparci |  |  |
| --- | --- | --- | --- | --- | --- | --- |
| Number of genomes | Sketch | Distances |  | Sketch (sketchlib) | Distances (sketchlib) | Distances (with prefilter) (sketchlib) |
| 10,000 | 1.17h | 34.8min | 58.8min<br>(30.9 CPU hours) | **36s**<br>(12.5 CPU minutes) | **1.08s** | 36s |
| 100,000 | 5.8h<br>(5.1 CPU hours) | DNF | DNF | **4.8min**<br>(1.7 CPU hours) | **1.3min**<br>(38.04 CPU minutes) | 6min<br>(101.21 CPU minutes) |
| 500,000 | - | - | DNF | **25.4min**<br>(8.3 CPU hours) | **0.5h**<br>(15.3 CPU hours) | **0.5h**<br>(9.82 CPU hours) |
| 1,000,000 | - | - | DNF | **48.0min**<br>(16.5 CPU hours) | 2.0h<br>(62.6 CPU hours) | **1.0h**<br>(22.7 CPU hours) |
| 5,200,000 | - | - | DNF | **4.8h**<br>(85.4 CPU hours) | 54.3h<br>(1,728 CPU hours) | **9.3h**<br>(213.8 CPU hours) |

### Network creation

Genome similarity networks are constructed from pairwise distances computed by the sketching algorithm. The distance scores are filtered using vectorised operations to retain only genome pairs with a similarity ≥98% (this is the default threshold, but can be changed by the user). Each genome is represented as a node in the network, with edges connecting genomes that exceed the similarity threshold. Genome similarity networks are constructed using NetworkX (Hagberg *et al*. 2008). The resulting connected components define our GCUs. To assess whether GCUs represent biologically meaningful groups, we taxonomically annotated each genome using GTDB-Tk (Chaumeil *et al*. 2022).

### Completeness correction

MAGs often exhibit genome incompleteness, which reduces similarity scores when compared with complete genomes from the same species (Shaw and Yu 2023). For example, if genomes 1 and 2 are identical and complete, they have a Jaccard similarity of 1. If genomes 1 and 2 are identical but genome 2 is only 80% complete (and only 80% of the k-mers match), they have a Jaccard similarity of 0.8. To address this systematic bias, we developed a completeness correction formula that adjusts the Jaccard similarity based on the estimated completeness of both genomes being compared. Define *c*_1_ as the completeness of genome 1, and *c*_2_ as the completeness of genome 2. The systematic reduction in |*S*1| and |*S*2| scale linearly with *c*_1_and *c*_2_, respectively, and the bias of |*S*1 ∩ *S*2| scales linearly with *c*_1_ *c*_2_because this expression depends on the completenesses of both genome 1 and 2, assuming these completenesses are uncorrelated. Hence, |*S*1 ∩ *S*2| can be corrected by dividing by *c*_1_. *c*_2_ and |*S*1| + |*S*2| − |*S*1 ∩ *S*2| can be corrected by dividing by *c*_1_ + *c*_2_ – *c*_1_ ⋅*c*_2_. This is equivalent to the following correction to the Jaccard index:

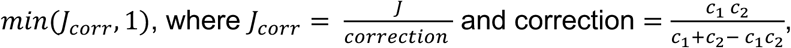

where J represents the observed Jaccard similarity, and c₁ and c₂ represent the completeness estimates for the two genomes. Genome completeness estimates (*c*_1_ and *c*_2_) are obtained from genome quality assessments such as CheckM (Parks *et al*. 2015) or CheckM2 (Chklovski *et al*. 2023), supplied by the user. If completeness values are not provided, the algorithm proceeds without completeness correction and assumes 100% completeness. We apply the completeness correction only to MAGs that are sufficiently complete to avoid overcorrecting the Jaccard distance, which could artificially inflate similarity scores to unrealistic values. As a default threshold for sufficient completeness, we chose *c*_1_ ⋅ *c*_2_ ≥ 0.64 (corresponding to an estimated completeness of ∼80% for both genomes). Below this threshold, corrections can exceed theoretical maximum values, potentially introducing false similarities (Supp. Figure 2). This conservative approach ensures that completeness correction is applied only to near-complete genome pairs.

To validate our completeness-correction algorithm, we compared clustering of the Unified Human Gastrointestinal Genome (UHGG) collection (Almeida *et al*. 2021) before and after correction. MAG completeness correction had a substantial impact: it reduced the total number of clusters by 35% (16,927 to 11,064) and singletons by 36% (9,919 to 6,369), thereby connecting thousands of MAGs to their true species clusters. To verify that completeness correction does not cause genomes of different species to be incorrectly grouped together, we assigned taxonomy as before using GTDB-Tk. We identified all 9,919 genomes that were singletons (single-genome clusters) before completeness correction and examined their cluster membership after correction. Of these, 3,550 (35.8%) joined an existing GCU. For every one of the 3,550 singletons that joined a cluster, we checked whether the genome’s assigned species matched the species of the GCU. In all 3,550 cases (100%), the singleton’s species matched the species already present in its new GCU. Critically, no genomes were incorrectly connected to wrong-species clusters, demonstrating that the correction accurately models the systematic bias of incompleteness without introducing false positives.

### Parameter configuration

For all analyses, a k-mer size of k=31 and sketch size s=1000 were used. To determine the optimal similarity threshold and knn (k-nearest neighbours) values for network construction, we performed a systematic sweep over similarity thresholds (0.95–0.99) and knn values (50–500) using a reference set of 218,859 high-quality (≥90% complete and less than 5% contaminated) genomes, which broadly cover the procaryotic space, with GTDB taxonomic annotations, evaluating clustering quality by purity and split species count. Across the tested range, the knn parameter had a negligible effect on clustering quality, while the similarity threshold was the primary driver of both purity and species fragmentation (Supp. Table 2). At the conventional ANI species cutoff of 0.95, cluster purity was only 94.2%, indicating that a substantial proportion of clusters contained genomes from more than one taxonomic lineage. Purity increased steadily with threshold, reaching 99.82% at 0.98, with approximately 2,757 split species. Beyond 0.98, species fragmentation increased sharply with little further gain in purity, indicating diminishing returns. We therefore selected a threshold of 0.98 as the optimal balance between avoiding spurious merging of distinct species and minimising unnecessary fragmentation of genuine ones.

### Refinement of clusters connected by chimeric genomes

Connected components within the similarity network are identified using NetworkX graph algorithms, with each component potentially representing a species cluster. However, chimeric genomes, in which two species have been erroneously assembled into a single genome, will create artefactual bridges between distinct genomic lineages. To address this, we apply a refinement algorithm based on network topology analysis. Contaminated genomes have characteristic signatures: they connect otherwise separate clusters (high betweenness centrality, as shortest paths between clusters pass through them), have few genuine similarities (low degree), and their neighbours are poorly connected to each other (low clustering coefficient).

We identify suspected bridge nodes as those with a high betweenness centrality. For that, we sort the betweenness values to find a large gap between consecutive values and return the value just after the gap. We then check whether the degree and clustering coefficient are below the median; nodes with this property are flagged as bridge nodes. Similarly, bridge edges connecting otherwise distinct components are removed entirely based on their extremely high edge betweenness values. This refinement ensures that final clusters reflect genuine genomic relationships rather than chimeric artefacts.

### Querying new genomes to assign GCUs

As an additional feature, we added a query mechanism to *gemsparcl* to enable any number of genomes to be assigned to a pre-existing GCUs from *gemsparcl*. For example, this allows taxonomically unknown MAGs to be rapidly taxonomically annotated by assigning them to GCUs. For querying, query genomes are sketched using the same k-mer length and sketch size as the reference database. K-mer length and sketch size are read in from the reference database to ensure comparability between databases. The distances are computed only between query genomes and the reference database (not all-vs-all), thereby making new genome assignment efficient by avoiding recomputation of the full reference distance matrix. Each query genome is assigned to a cluster if it has at least one reference genome with an ANI above the ANI threshold (0.98); the cluster of the best-matching reference genome determines the GCU assignment. If a query genome hits reference genomes from multiple non-singleton clusters, it is flagged as potential contamination (but could be a genuine, high-quality, bridging genome and warrants further analysis). If it hits both a non-singleton cluster and a singleton, it is flagged as “connecting_cluster_and_singleton”. If no reference genome exceeds the threshold, the query genome is left unassigned (no hit), indicating a potentially novel species not represented in the reference database.

### The dataset encompasses 5.6 million bacterial genomes

To assess scalability and biological accuracy at scale, we constructed a dataset of 5,645,186 genomes (≥80% complete and less than 5% contaminated) integrating five large public repositories, including both culture-based isolates and MAG assemblies from diverse environments: AllTheBacteria (2.4M isolates), RefSeq (465K isolates), MGnify (360K MAGs and isolates), SPIRE (604K MAGs), and mOTUs (1.8M MAGs).

### Tree methods for investigating genomes from the International Space Station

The *Pantoea piersonii* phylogenetic tree was constructed from 66 non-redundant genomes (drawn from two *gemsparcl* GCUs, with 13 duplicates removed) annotated using Prokka (Seemann 2014). A core-genome alignment was generated with Panaroo (Tonkin-Hill *et al*. 2020) and a maximum-likelihood tree inferred using VeryFastTree (GTR + gamma model). The pangenome was characterised using CELEBRIMBOR (Hellewell *et al*. 2024) with core-corrected genome frequency thresholds to account for genome incompleteness in MAGs.

### Tree methods for placing unknown MAG cluster phylogenetically

To place *UBA11524 sp000437595* (from the 5.6 million genome dataset) phylogenetically, we first identified its closest relatives by querying the ten most complete genomes from this GCU (cluster 45) against all 5.6 million genomes in our dataset using sketchlib. The 10 GCUs with the highest median ANI to the cluster 45 representatives were selected, and the most complete genome from each was extracted as a representative. Together with the cluster 45 MAG itself, this yielded a set of 11 genomes with a median ANI range of 79.9-84.3% relative to the focal species. To construct a phylogenetic tree, we used GTDB-Tk v2.4.1 (Chaumeil *et al*. 2022) with GTDB release 226 to identify and align the 120 bacterial marker genes (bac120), and inferred a maximum-likelihood phylogeny from the concatenated alignment using IQ-TREE v3.0.1 (Nguyen *et al*. 2015) under the LG+G4 substitution model with 1,000 ultrafast bootstrap replicates. The resulting tree was midpoint-rooted for rendering.

## Results

### *Gemsparcl* clusters over five million bacterial genomes under 24 hours

We benchmarked the performance of *gemsparcl* in two stages: first, by comparing the underlying sketchlib.rust library against established ANI estimation tools (Mash (Ondov *et al*. 2016), sourmash (Pierce *et al*. 2019), and skani (Shaw and Yu 2023)), then evaluating the complete *gemsparcl’s* genome clustering workflows against dRep’s genome clustering workflows. We first evaluated sketchlib (v0.2.4), sourmash (v4.8.14), and skani (v0.3.0) across four dataset sizes (10,000, 100,000, 500,000, and 1 million genomes), performing all-versus-all pairwise distance calculations, utilising 32 threads. Sketchlib demonstrated the best performance across all dataset sizes (Table 1, Figure 2). The performance difference becomes particularly pronounced with large datasets. For the 1 million genome dataset, sketchlib completed sketching and distance calculations in just under two hours, compared with at least 58 days (extrapolated) for sourmash and 130 days (extrapolated) for skani. This corresponds to a 762× speedup over sourmash and a 1,700× speedup over skani.

To evaluate performance at scale, we benchmarked the runtime of our implementation on 5.6 million genomes, representing trillions of pairwise distances. While sketching-based approaches are faster than alignment-based methods, they still suffer from the same all-vs-all quadratic growth. For example, pairwise comparison of 1,000 genomes results in 1×10^6^ comparisons, while comparison of 1,000,000 genomes results in 1×10^12^ comparisons; a number too large to feasibly be processed by even the fastest of existing genome clustering tools. By using an inverted index, *gemsparcl* reduces this complexity (Supplementary figure 3) and can cluster 5.6 million genomes using 48 threads, taking around 4.5h for sketching, less than 3h for inverted-index calculations, and around 5h for distance calculations. Network calculation took about 1.5 hours, totalling ∼14 hours and using no more than 16.5 GB of memory.

We subsampled 5,000 genomes for comparison with dRep. As shown in Table 2, *gemsparcl* demonstrates a substantially faster runtime than dRep. For a very small dataset of 5,000 genomes, dRep requires 7.5 CPU hours while *gemsparcl* requires only 0.1 CPU hours, a 532× speedup. This performance gap widens with larger datasets: dRep becomes computationally infeasible for datasets in the range of tens of thousands of genomes (which already require days of compute time), whereas *gemsparcl* scales efficiently to such datasets.

**Table 2.**
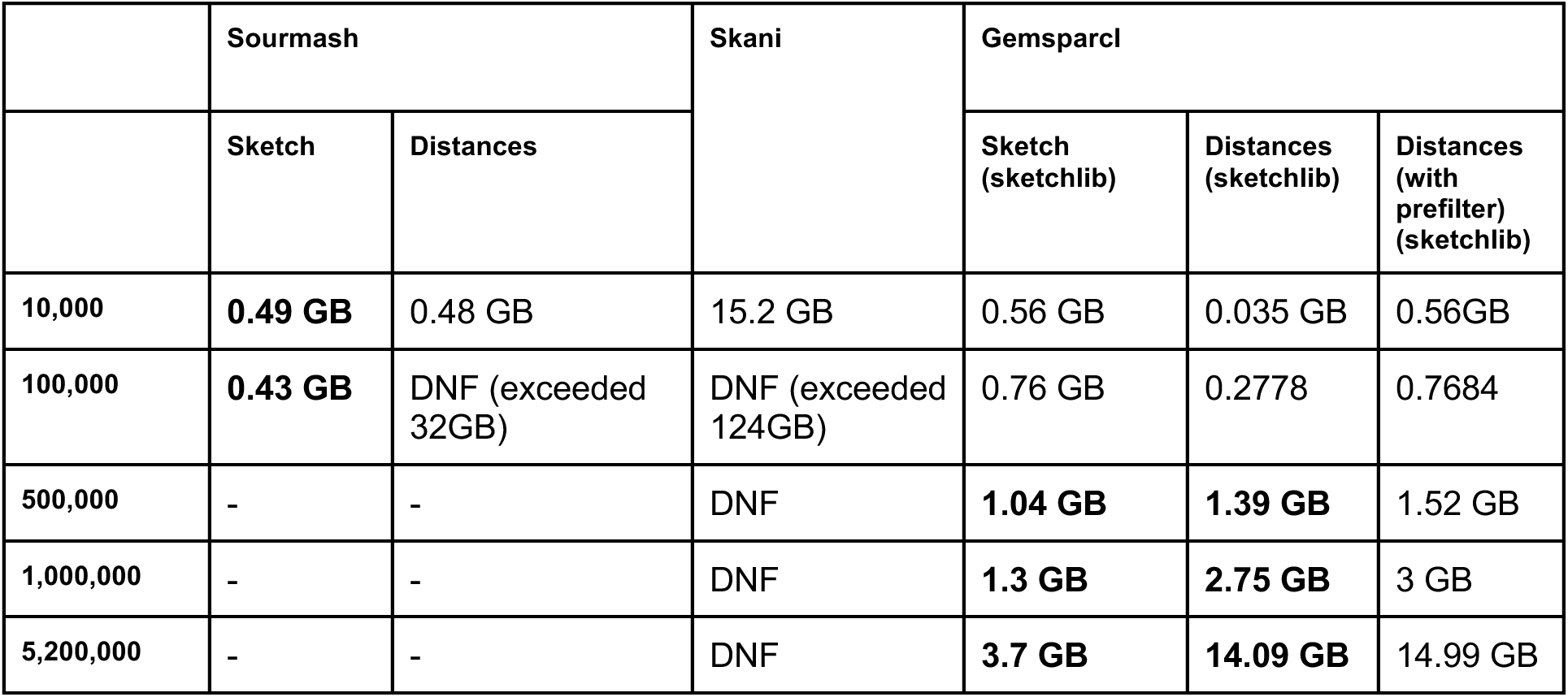
Peak memory usage of pairwise genome similarity tools across dataset scales. Benchmarking was performed on an Intel Xeon Gold 6336Y processor (2.40–3.60 GHz, 32 threads, and up to 124 GB RAM). DNF indicates the tool exceeded available system memory before completing. Dashes indicate tools that could not be evaluated at that scale due to prior failure on a smaller dataset.

### Plasmid content does not affect clustering accuracy

We investigated the impact of plasmids on ANI estimation, as plasmids might lead to larger accessory distances at short evolutionary times. To assess whether plasmid sequences impact clustering results, we analysed genomes from two prevalent human gut species: *Bacteroides uniformis* and *Phocaeicola vulgatus*. We used high-quality isolate genomes from 59 *B. uniformis* isolates and 48 *P. vulgatus* isolates. These genomes were sequenced using PacBio’s long-read technology, resulting in highly contiguous assemblies, with most genomes assembled into a single circular contig (ranging from 1 to 7 contigs, including 40 genomes assembled into a single contig). The plasmids were derived from smaller circular contigs, and most contained the repA protein. We compared clustering results before and after removing all known plasmids. Plasmid number and size varied across genomes, ranging from 6,993 bp (0.13% of genome size) to 134,896 bp (2.55% of genome size).

Plasmid removal altered some pairwise ANI values but did not change the overall clustering structure. Genomes that clustered together remained in the same clusters. Comparing pairwise ANI scores before and after plasmid removal (Supplementary Figure 6) revealed minimal changes in the similarity range relevant for clustering. For genome pairs with an ANI above 98% (our clustering threshold), the data points aligned tightly with the perfect correlation line, indicating that plasmid content had a negligible impact on similarity estimates. Lower-similarity genome pairs (<98% ANI) exhibited greater variation. However, these comparisons fall below our clustering threshold and therefore do not influence cluster assignment. We were able to establish that no data moved to the wrong side of the decision boundary.

### Querying validation

To validate that the querying works as expected, we used 316,238 genomes from MGnify (quality-filtered to pass the QS50 score (Parks *et al*. 2022) with varying genome completeness statistics (mean completeness: 85.9%; mean contamination: 1.21%). First, as the validation set, we clustered all genomes to obtain their cluster assignments. Then, one representative genome was held out from each of the 200 largest clusters to provide the ground-truth assignment, and the remaining 316,038 genomes were re-clustered to form the reduced, hold-out-free reference database. The 200 held-out genomes were then queried against this new reference database.

All 200 held-out genomes were correctly reassigned to their original clusters. Notably, four cases highlighted the structural importance of individual genomes to cluster connectivity. Specifically, in two cases, the held-out genome was the sole connection between a fringe genome and its cluster, so removing it made the fringe genome a singleton. Querying the held-out genome restored the connection. In the other two cases, the held-out genome had been bridging two sparsely connected subclusters; removing the hold-out genome split the originally sparsely connected cluster into separate clusters. Again, the querying of the holdout genomes reconnected the split clusters.

### *Gemsparcl* creates genomically coherent units which enhance taxonomical classifications

The dataset encompassing 5.6 million bacterial genomes allowed us to evaluate whether species-level resolution is maintained across genomes of varying quality and origin, and to assess how culture-based and metagenomic sampling strategies capture overlapping or complementary bacterial diversity. In our method, we have balanced two opposing failure modes: over-clustering, where the threshold is overly permissive and merges distinct species into the same GCU, resulting in a multi-species cluster, and under-clustering, where the threshold is too strict and splits a single species across multiple GCUs, as well as inflating the number of singletons. Singletons, genomes with no neighbours above the ANI threshold that therefore form no cluster, are a natural outcome for genuinely rare or divergent lineages, but an excess suggests the threshold is too stringent. Evaluating both GCU purity and singleton numbers is therefore necessary to distinguish a well-calibrated threshold from one that is too strict in either direction.

Clustering of the 5.6M dataset identified 92,954 GCUs and 101,656 singletons (of the singletons, 76,563 are MAGs), demonstrating scalability to over 5.6 million genomes. Of the 92,954 GCUs, only 221 (0.24%) contain more than one GTDB species label, giving a purity score of 99.76%. Inspecting the genomic composition of these clusters further, the multi-species label clusters fall into one of two categories: 93 GCUs where genomes share a genus but span multiple unnamed species (sp* identifiers), as seen in Figure 3, and 128 where two or more formally named species co-occur within the same genus. The small fraction of impure GCUs likely reflects the biological reality that ANI-based species boundaries are not always discrete. Parks et al. (Parks *et al*. 2022) demonstrated that approximately 2.2% of GTDB species clusters are’fuzzy’, containing genomes with interspecific ANI values ≥95% that blur species boundaries. The 221 mixed GCUs are therefore expected, rather than an inherent methodological issue, and arise because some closely related species simply do not have a clear ANI boundary between them. GTDB resolves this by anchoring species to type strains, but when clustering based solely on sequence similarity, these species are difficult to distinguish.

**Figure 3:**
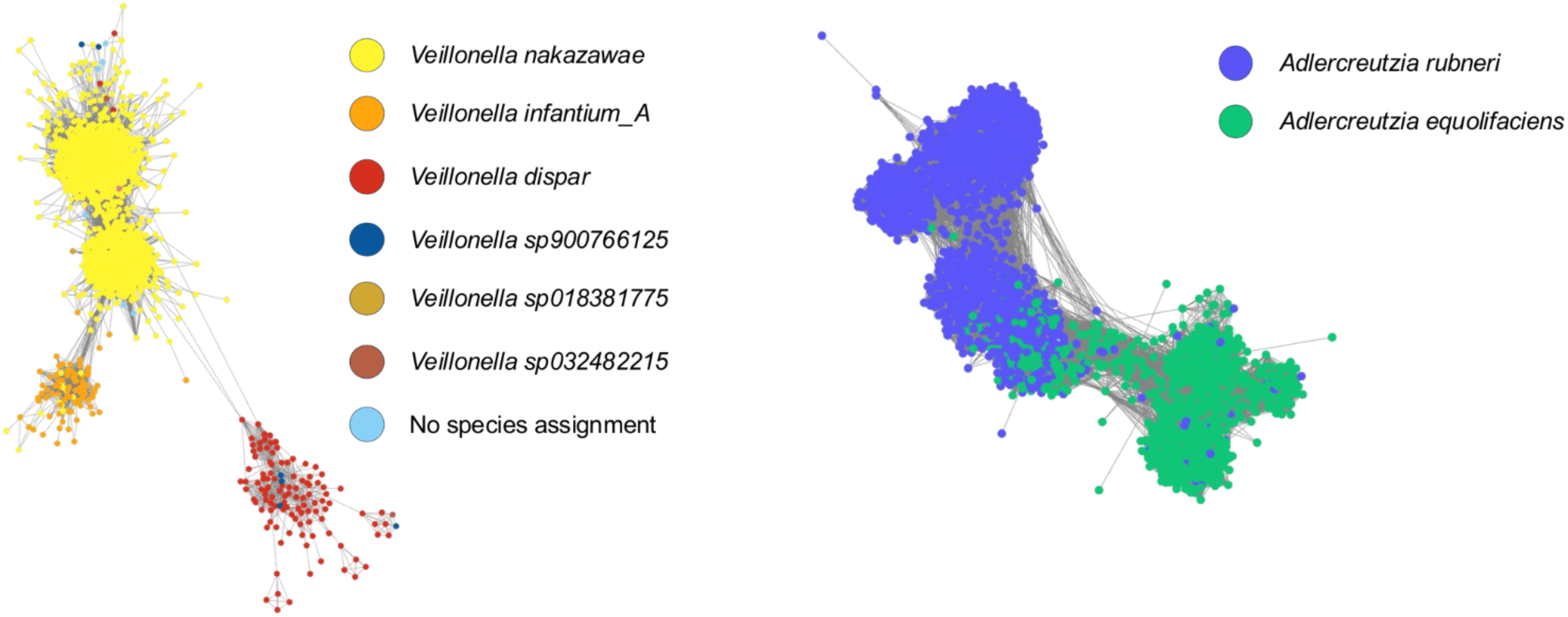
Fuzzy species clusters: Genomes with different taxonomic assignments clustering together in the same GCU. Left: seven different *Veillonella sp.* are brought together, with *V. dispar, V. nakazawae and V. infantium_A* largely forming distinct sub-clusters. Right: two *Adlercreutzia* species that do not display any strong separation from each other. These are two examples of taxonomic species that are indistinguishable by this approach, which clusters everything together.

We investigated the distribution of culture-based and metagenomic sampling across major species. Of the 92,954 GCU, 10,163 GCUs (5.08%) contained both isolates and MAGs, yet this accounts for 73.28% of all clustered genomes (4,062,097 genomes). The three largest GCUs of *Salmonella enterica* (639,314 genomes: >99.9% isolates), *Escherichia coli* (481,406 genomes: 93.8% isolates), and *Mycobacterium tuberculosis* (190,789 genomes: 100% isolates) demonstrated that while both data types cluster together when present, major pathogen species remain predominantly sampled through culture-based approaches. Examining the 25 largest GCUs revealed that they were dominated by isolate genomes of major pathogens, which accounted for 88.1% of the genomes, reflecting both widespread culture-based sequencing efforts for these pathogens and the rarity of these pathogens in datasets from environmental samples. In contrast, MAG datasets contributed primarily to species-level diversity, with MAGs derived from human gut metagenomic samples contributing to 5,450 distinct GCUs and marine MAGs contributing to 8,233 GCUs (including 63 and 19 multi-species clusters, respectively).

We further investigated the 221 GCUs that contained more than one taxonomic label. Two examples of GCUs which are taxonomically labelled as different species but genomically indistinguishable.

An example of an impure cluster is GCU 155, where *Bacillus subtilis* and *Bacillus licheniformis* had been incorrectly merged by two taxonomically unannotated bridge nodes; after refinement, GCU 155 was resolved into two distinct clusters (Supplementary Figure 5). In total, 17 GCUs required refinement, and in each case where a bridge node was removed, that genome could not be assigned a species-level annotation by GTDB-Tk.

To quantify over-clustering, we assigned each non-singleton GCU its species identity based on its GTDB annotation. Of the 92,954 GCUs, 65,407 carried a species-level label, representing 48,169 distinct species (Supp. Table 4). Of these, 42,461 species (88.2%) were represented by a single GCU, while 5,708 species (11.8%) were fragmented across two or more GCUs, resulting in 17,238 excess clusters. Importantly, the vast majority of these additional clusters were very small: 90.3% contained just 2-9 genomes, 8.1% contained 10-99 genomes, and only 1.7% reached 100 genomes or more. This indicates that over-clustering is predominantly a boundary effect, where a small number of genomes within a species have pairwise ANI values that fall marginally below the 98% threshold, causing them to split off into a separate cluster rather than join the main species GCU. This is an expected consequence of applying a discrete similarity cutoff to a continuous distribution of ANI values, and does not reflect a systematic failure to group genuinely divergent populations. Among fragmented species, 3,673 (64.4%) were split into exactly two GCUs, further supporting that most cases represent marginal splits near the threshold rather than severe biological fragmentation.

Two distinct patterns of over-clustering emerge. The first involves heavily sequenced species where one GCU captures the vast majority of genomes, but the species additionally fragments into several large satellite GCUs. This pattern is driven by high within-species diversity that partially exceeds the 98% ANI threshold. For example, 96.29% of the 663,941 *Salmonella enterica* genomes reside in a single GCU, yet the species spans 42 large GCUs (≥100 genomes) in total, reflecting its exceptional diversity across more than 2,500 described serotypes (Gorski *et al*. 2022). Similarly, *Escherichia coli* (484,874 genomes; 99.28% in the largest GCU), *Streptococcus pneumoniae* (131,730 genomes; 92.27% in the largest GCU), and *Campylobacter jejuni* (99,708 genomes; 90.42% in the largest GCU) span 11, 27, and 19 large GCUs respectively, each representing intensively sequenced clinical pathogens with well-documented within-species diversity (C. B. Blacklock *et al*. 2024; Siniagina *et al*. 2021).

The second pattern is near-complete fragmentation with no dominant GCU, seen in species with exceptionally high genomic diversity. The most striking example is *Helicobacter pylori*, where 4,833 genomes are distributed across 619 GCUs and 1,002 singletons, with the largest GCU containing only 456 genomes (9.44% of all *H. pylori* genomes). This reflects the exceptional recombination rate and mutational diversity of *H. pylori*, with distinct strains known to co-occur at different sites within the same human gut (Blaser and Berg 2001; Wilkinson *et al*. 2022). A similar pattern is seen in the *Collinsella* genus, where 26,668 genomes cluster into 7,728 GCUs, with the largest containing only 1,672 genomes, consistent with the high phylogenetic diversity within this genus. In both cases, over-clustering reflects genuine biological diversity rather than a methodological limitation of the 98% ANI threshold.

Singletons, as an additional manifestation of over-clustering, can arise either from genuine biological rarity and diversity or from low assembly quality. To distinguish between these, we first validated singleton quality. CheckM analysis of MAG singletons revealed a mean completeness of 90.7% (±6.0%) and contamination of 1.6% (±1.4%), comparable to MAGs in GCUs (92.6% complete, 1.6% contaminated), confirming that singletons are not predominantly low-quality assemblies. Examining the taxonomic composition of singletons further supports the notion that singletons represent genuine diversity. Of 101,656 singletons, 53,510 (52.6%) carried a species-level annotation, 41,844 (41.2%) were assigned only at genus level, and 6,302 were assigned above the genus level by GTDB. Among species-level singletons, 28,515 (55.8%) were species with no matching genomes in the clustered dataset, including 24,572 unique species seen exclusively as singletons, 91.8% of which were represented by a single genome. This strongly suggests these taxa are genuinely rare rather than clustering artefacts.

Singleton rates (Supp. Table 3) varied substantially by genome source and environment, consistent with this interpretation. Isolate genomes showed a 0.87% singleton rate compared to 2.76% for MAGs, reflecting both the greater sampling depth of culture-based sequencing and the higher biological diversity captured by metagenomic approaches. Environmental datasets showed the highest singleton rates, with soil (72%), tomato rhizosphere (49.79%), and zebrafish faecal (40.24%) environments having the largest proportion of genomes with no close relatives in the full 5.6 million genome dataset, pointing to substantial uncharacterised diversity in these undersampled niches. In contrast, the human gut showed only 0.71% singletons, consistent with the frequent sampling this microbiome has received.

**Table 3.** Computational performance comparison of clustering tools. Runtime comparison of *gemsparcl* and dRep on 5,000 genomes using 32 threads. *Gemsparcl* demonstrates substantially reduced processing time (532 times faster) compared to dRep (v3.2.2) run in dereplication mode with a 95% ANI threshold and no genome quality filtering using the default clustering algorithm ANImf (Richter and Rosselló-Móra 2009).

| 5,000 genomes | dRep | Gemsparci |  |
| --- | --- | --- | --- |
|  |  | Normal | Inverted (with prefilter) |
| Time | 4.14h (7.5 CPU hours) | 33 s (0.1 CPU hours) | 28s (0.2 CPU hours) |
| Memory | 3.61 GB | 0.51 GB | 0.51 GB |

Overall, elevated singleton rates are best explained by genuine high genomic diversity rather than methodological limitations, and are most pronounced in environments where bacterial diversity remains poorly characterised. *Gemsparcl*, in combination with taxonomic labelling, provides a means to both discover and organise this diversity at scale, with cluster representatives offering a practical entry point for downstream exploration of understudied lineages.

### *Gemsparcl* enables universal analysis of isolate and metagenome data

Within the full dataset, there are many true duplicate genomes. For example, for isolate genomes, these may arise from the sequencing of the same type strain or epidemiological sampling campaigns where the same organism has been transmitted between two individuals. Within the MAG collections, there is also redundancy between the datasets, due to: (i) the assembly of the same underlying sequence dataset being used to generate MAGs; (ii) replicates (technical and experimental), both leading to the reconstruction of the same, or near identical, genome; and (iii) genomes being shared across different samples. To remove completely redundant genomes, we randomly removed one genome from each pair with 100% ANI similarity as determined by *gemsparcl*. Starting with 5,657,541 genomes, we first removed 655,541 genomes from MAG-MAG pairs, then 13,913 genomes from MAG-isolate pairs (preferentially keeping isolates), and finally 1,500,869 genomes from isolate-isolate pairs. This resulted in a final non-redundant dataset of 3,487,218 genomes (Figure 4). It highlights that MAG reconstruction can generate genomes equivalent to isolates, and that the most redundancy exists among isolate genomes.

**Figure 4:**
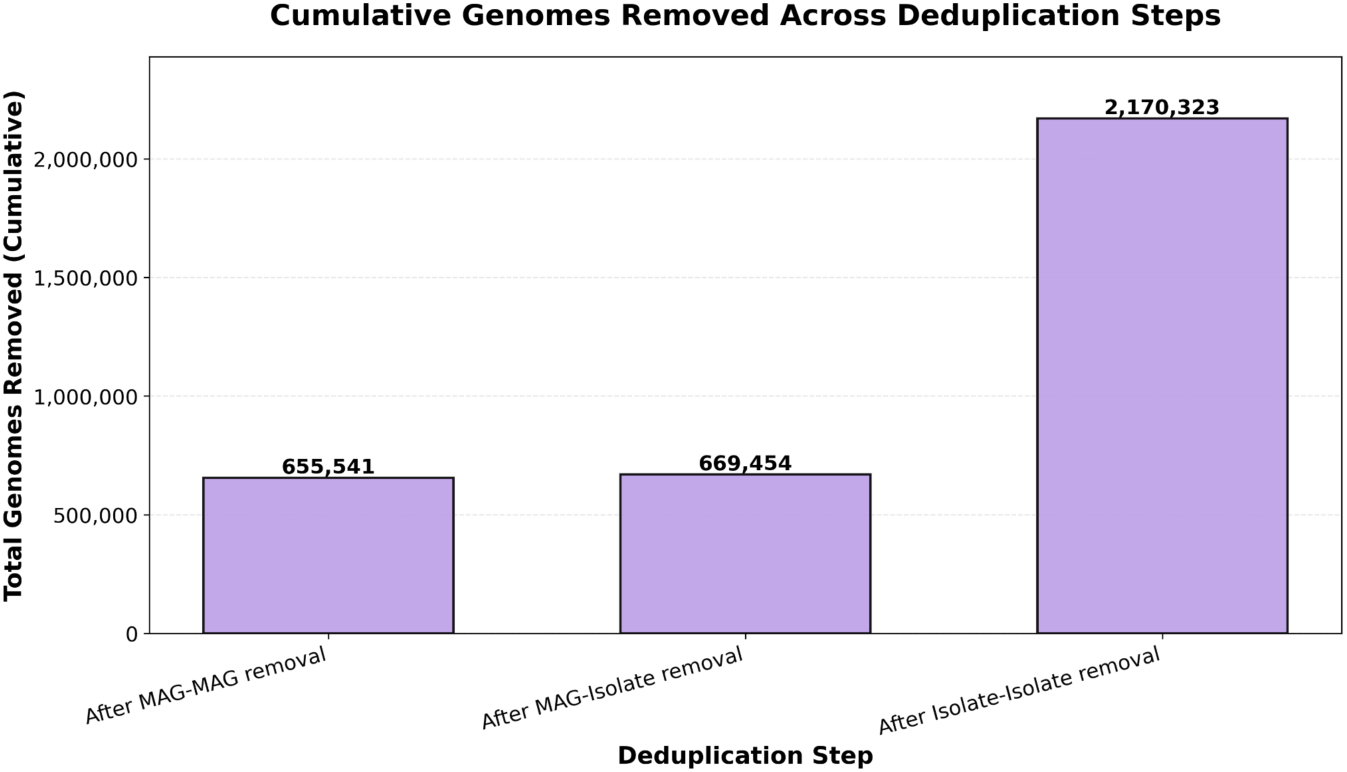
Plot showing redundancy removal of genomes. Starting with 5,657,541 genomes, we first removed 655,541 genomes from identical MAG-MAG pairs, then 13,913 genomes from identical MAG-isolate pairs (preferentially keeping isolates), and finally 1,500,869 genomes from identical isolate-isolate pairs.

Investigating the GCU composition across datasets, especially isolates and MAGs, it was evident that most GCUs are either MAG- or isolate-dominated (Figure 5). Beyond characterising known diversity, *gemsparcl* enables the identification of novelty within large genome collections. GCUs that lack isolate representatives and carry no formal species annotation represent candidates for undescribed species. These have been detected through metagenomic sequencing but never successfully cultured. To identify the most prevalent of these, we searched for the largest MAG-only GCUs lacking isolate representatives, which could be important targets for developing isolation and culturing methods. The largest such GCU corresponds to the species *UBA11524 sp000437595* (UBA = Uncultivated Bacteria and Archaea), which contains 10,504 genomes after duplicate removal. This species belongs to the genus *UBA11524* within the family *Aristaeellaceae,* and, based on current literature and available sample metadata, it is prevalent in mammalian gut environments, including humans, but little is currently known about the role it plays (Supp. Figure 4). Geographically, *UBA11524 sp000437595* appears to have a global distribution, with the majority of samples originating from Europe, followed by Asia and North America, which broadly reflects the distribution of microbiome sequencing efforts.

**Figure 5:**
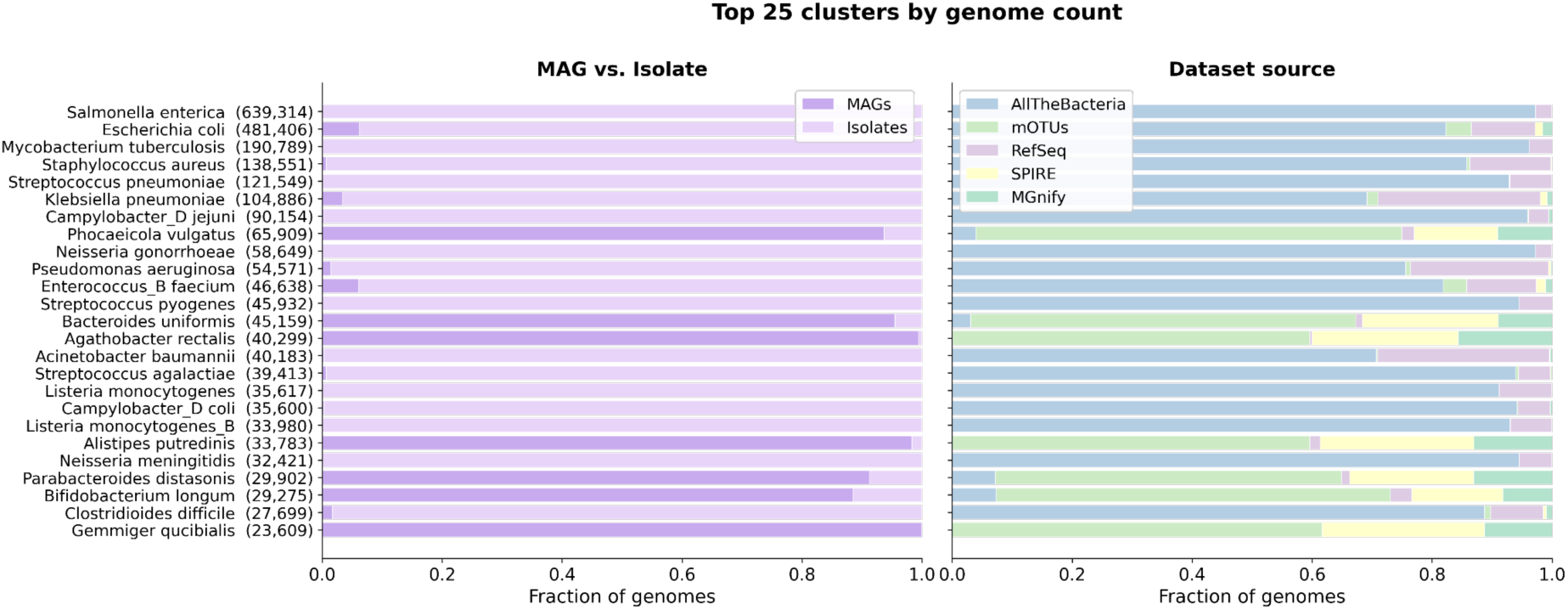
Composition of the 25 largest GCUs and their source datasets. (Left) Proportion of MAG and isolate genomes within each of the 25 largest GCUs, with total cluster size shown in parentheses. (Right) Genome composition by source dataset across the full 5.6 million genome database.

**Figure 6:**
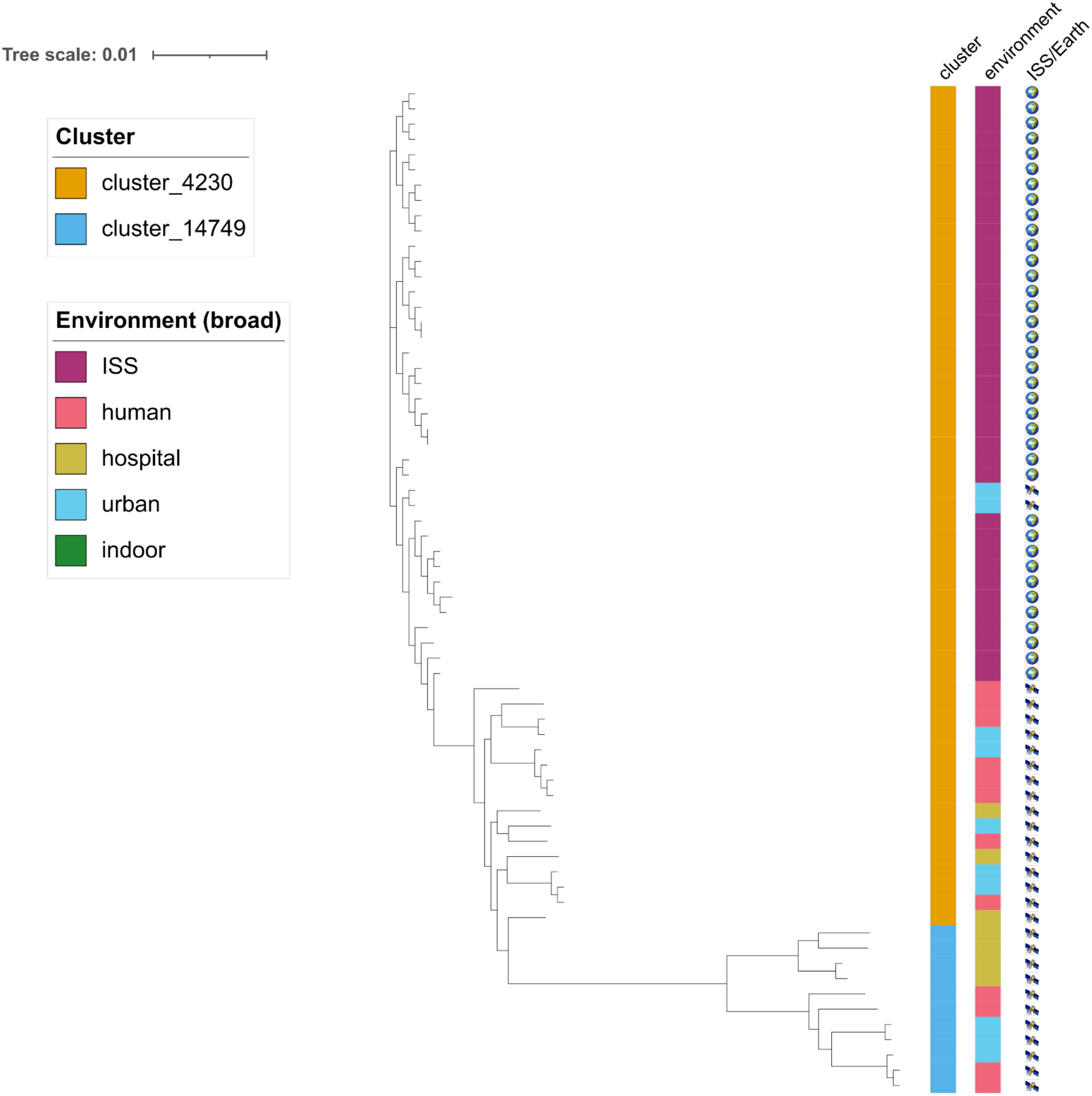
Core genome phylogeny of 66 *P. piersonii* genomes from GCU 4230 and GCU 14749, inferred from a Panaroo core genome alignment using FastTree. Space station and terrestrial genomes form distinct clades, with GCU 14749 (terrestrial) positioned on a long branch indicating substantial divergence from GCU 4230.

To further investigate this species, we performed a pangenome analysis using CELEBRIMBOR (Hellewell *et al*. 2024). Despite all genomes clustering together at ≥ 98% ANI, the species shows considerable genomic diversity. The core genome, defined as the set of genes present in ≥92.65% of genomes, after accounting for incomplete MAGs, consists of 1,019 genes. The intermediate accessory genome comprises 2,235 genes, while the rare shell genome, present in ≤10.56% of genomes, is remarkably large at 175,481 genes, suggesting an open pangenome with extensive strain-level variation.

To investigate the phylogenetic context of this GCU, we queried the ten most complete representative genomes against all 5.6 million genomes in the database using sketchlib. The closest match was Vescimonas sp000435555 at 88.1% ANI, well below both the 95% ANI threshold commonly used to delineate bacterial species and the 98% threshold used for clustering in this study. Notably, even these nearest neighbours are not genuinely close relatives; at 88.1% ANI, the best hit shares neither genus nor family with this GCU, with the closest shared taxonomic rank being order. This means that despite searching across 5.6 million genomes, no close relatives of this lineage exist in any currently catalogued bacterial genome database. The ten nearest neighbour GCUs span three phyla (*Bacillota_A*, *Bacteroidota*, and *Actinomycetota*), with no neighbour sharing the same family (*Aristaeellaceae*) and only one (*Limadaptatus sp900554085*) belonging to the same order (*Christensenellales*). This phylogenetic isolation, combined with its abundance in mammalian gut environments, highlights *UBA11524 sp000437595* as a high-priority target for cultivation and characterisation.

A related powerful application of *gemsparcl* is the ability to search for closely related genomes across large-scale bacterial genome databases. As an example, we used *Pantoea piersonii,* which was first isolated from the International Space Station (ISS), but its original terrestrial reservoir remains unclear (Crosby *et al*. 2023). Survival on the International Space Station requires bacteria to withstand microgravity, radiation, desiccation, and nutrient limitation, often by forming protective aggregates or entering stress-tolerant states (Koehle *et al*. 2023). To investigate the presence of similar *P. piersonii* genomes not coming from the ISS, we clustered 6 *Pantoea piersonii* isolates recovered from the ISS against all 5.6 million genomes. All six space station genomes clustered with other *P. piersonii* genomes (n = 55, GCU 4230) drawn from multiple independent datasets. We also identified a second *P. piersonii* GCU (n = 11, GCU 14749) containing no ISS genomes, with all members originating from terrestrial sources.

Pangenome analysis using CELEBRIMBOR revealed 11,620 gene clusters across all genomes from both *P. piersonii* GCUs. The core genome, defined as genes present in ≥93.85% of genomes (CGT-corrected), comprises 3,323 conserved genes, with a further 1,413 genes in the intermediate accessory genome. The rare shell genome, present in ≤10.77% of genomes (CGT-corrected), comprises 6,884 gene clusters, approximately 60% of the total pangenome. Examining the pangenome at the level of individual GCUs reveals striking differences between the two groups. Using the same thresholds as above, the shared core genome between the two GCUs comprises only 1,952 genes. Within each GCU, the core genome differs dramatically: only 49 genes pass the core threshold in GCU 14749, compared to 1,515 genes in GCU 4230. *P. piersonii* genomes from the ISS carried 16 mobile genetic elements at high frequency, including prophage integrases, insertion sequences from six families, and a transposon, which were rare or absent in terrestrial genomes from both phylogenetic clusters. Five of these elements were found exclusively in space station genomes and were completely absent from all terrestrial isolates, indicating that the accumulation of mobile genetic elements in this species is associated with the space environment. However, we cannot rule out a founder effect, whereby a single lineage carrying mobile genetic elements was introduced to the ISS and subsequently expanded, rather than reflecting ongoing acquisition driven by the space environment itself.

To visualise the genetic relatedness of genomes across both clusters, we annotated all genomes using Prokka (Seemann 2014) and constructed a core genome alignment with Panaroo (Tonkin-Hill *et al*. 2020), from which we inferred a phylogenetic tree using FastTree. The resulting tree shows a clear distinction between space station and terrestrial genomes. GCU 14749 is positioned on a notably long branch, indicating substantial divergence from all genomes in GCU 4230.

The ISS genomes form a cohesive cluster with a well-defined core genome, while the five mobile genetic elements found exclusively in space station isolates point toward ongoing genomic remodelling under the selective pressures unique to the ISS environment, including microgravity, radiation, desiccation and nutrient limitation (Koehle *et al*. 2023). Whether *P. piersonii* was introduced to the ISS from a terrestrial reservoir represented by GCU 14749, or from an as-yet-unsampled source, remains an open question that expanded environmental sampling could address. The genomic signatures identified here are more consistent with post-arrival adaptation to the ISS environment than with preadaptation.

### GCU cluster structure can represent subspecies structure

Finally, we investigated whether the clusters in *gemsparcl* capture meaningful genomic structure at the subspecies level. We extracted all *Staphylococcus aureus* genomes from the dataset, totalling 141,335. Examining the clustering structure revealed a visible substructure within this species cluster. As shown in Figure 7, the subclusters within this GCU align well with known clonal complexes typically used to define subspecies within *S. aureus*. Each clonal complex resolves to its own subcluster, including three subclusters without any clonal complex assignment.

**Figure 7:**
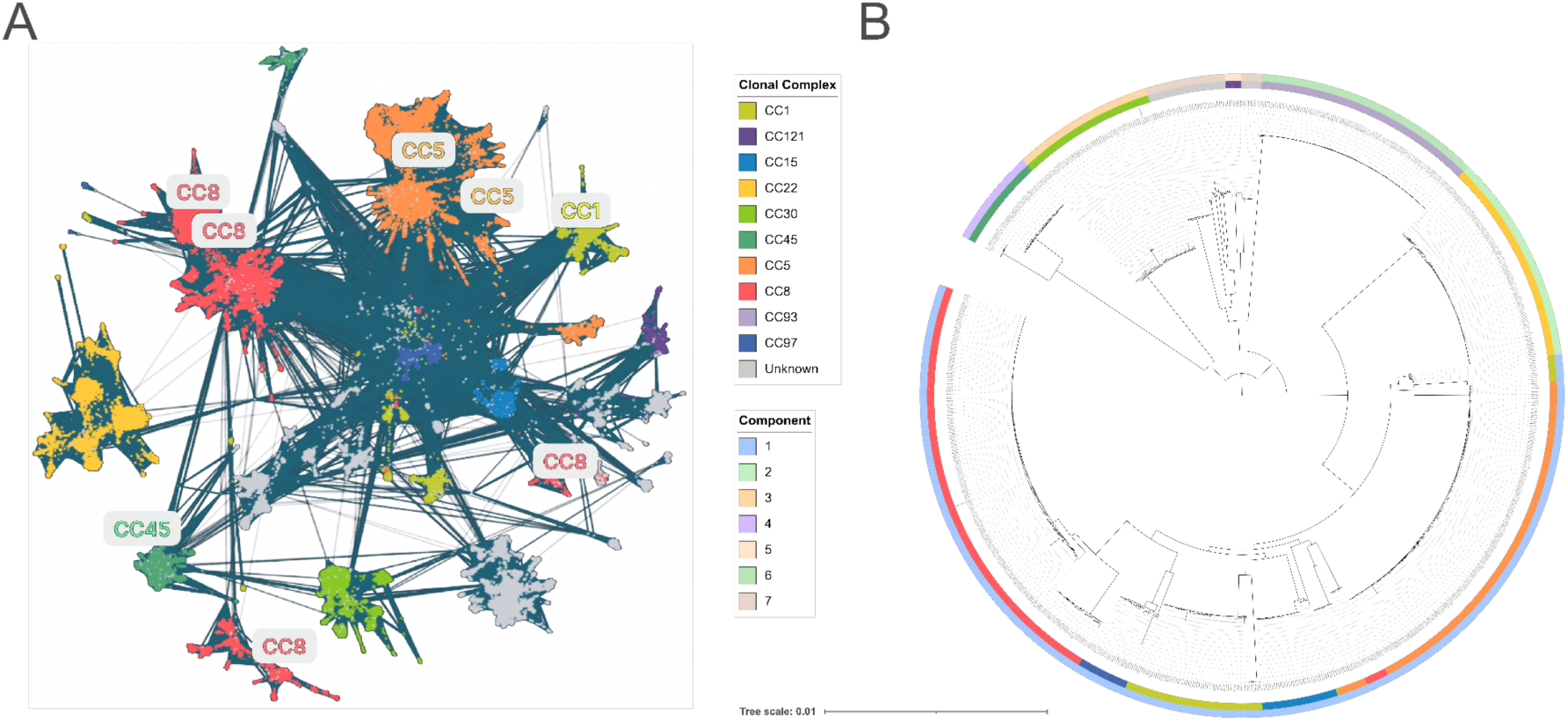
Subspecies structure within S. aureus revealed by *gemsparcl* clustering. (A) Substructure within the *S. aureus* GCU becomes clearly visible when visulising the network of genomes. (B) Core genome phylogeny of 1,000 *S. aureus* genomes, with the inner band coloured by clonal complex assignment and the outer band coloured by *gemsparcl* cluster identity, demonstrating concordance between phylogenetic groupings and cluster assignments.

A number of genomes cluster centrally and cannot be resolved into a distinct subcluster. This demonstrates that the single-linkage thresholds used in *gemsparcl* have limitations in subspecies resolution. Because we use a simple k-mer distance measure that incorporates both core and accessory genome components, this prevents precise identification of subspecies or sequence types. In *S. aureus*, core genome evolution is relatively clock-like (i.e., accumulating mutations at a roughly constant rate over time), but the genome contains numerous mobile genetic elements. A core genome phylogeny yields strongly separated clades (typically clonal complexes), but within these, there is substantial accessory variation due to mobile genetic elements (Enright *et al*. 2002). Achieving higher resolution will require methods based on variable-length k-mers, such as PopPUNK or other approaches better suited to capturing fine-scale genomic variation within species.

## Discussion

Reductions in sequencing costs have driven an exponential expansion of bacterial genome databases, with isolate collections and MAG repositories now each numbering in the millions. This growth has exposed a fundamental mismatch: traditional clustering tools scale quadratically with dataset size, making comprehensive database-wide analysis computationally intractable. Compounding this, the incomplete nature of MAGs has historically excluded them from large-scale comparative analyses, limiting our view of bacterial diversity to the culturable fraction. Until now, clustering millions of MAGs and isolates together in a single analysis has been effectively impossible within practical time and memory constraints.

We clustered over 5.6 million bacterial genomes into 92,954 GCUs in less than 14 hours using 48 threads and 16.5 GB of RAM. This represents a fundamental shift: comprehensive database-wide genome clustering is now computationally possible on standard hardware. For comparison, traditional all-versus-all methods such as dRep scale quadratically with dataset size, and would require months of computation and thousands of GB of RAM to cluster a dataset of this scale, making such analyses practically infeasible for most researchers. *Gemsparcl*’s improved scaling enables routine database clustering and real-time integration of new genomes, transforming a near-impossible task into standard practice.

Three technical innovations enable this performance while maintaining biological accuracy. First, binned sketching with an inverted index dramatically accelerates all-versus-all comparisons by restricting distance calculations to genomes that share k-mer content. Our inverted index approach reduces redundant comparisons, lowering the computational complexity. Second, explicit completeness correction accounts for MAG incompleteness, enabling accurate distance estimation between environmental genomes and high-quality isolate genomes. This correction is essential for integrating the growing volume of metagenomic data with traditional culture-based sequencing, ensuring that incomplete assemblies do not artificially deflate genomic distances. Third, k-nearest neighbour filtering improves computational efficiency. By retaining only the 50 most similar genomes for each query and applying a 98% similarity threshold, the method focuses edge creation on biologically relevant comparisons. This sparse network construction reduces the number of edges to be evaluated and stored, enabling efficient clustering of millions of genomes with an unbalanced taxonomic distribution while ensuring that connected components represent genuine genomic similarity.

The scalability of *gemsparcl* enables previously complex and/or computationally demanding analyses of microbial diversity and evolution to be performed on a typical single multi-core CPU. For example, microbiome studies often require comparing newly generated genomes against a reference set of isolates and/or MAGs. However, these reference datasets may be limited in the number of genomes, making it difficult to assess whether a genome represents a genuinely novel lineage or is simply absent from the reference database. By clustering all genomes derived from all available environments simultaneously, *gemsparcl* enables the detection of novelty in a true One Health context. A genome from a human gut study can now be directly compared against soil, animal, and marine datasets, revealing whether the species is shared across ecosystems and, when it is, how genomic variation is distributed across hosts, body sites, and ecosystems at an unprecedented scale.

For example, we demonstrated that *Pantoea piersonii* genomes from the International Space Station formed a distinct genomic cluster from terrestrial isolates, with five mobile genetic elements found exclusively in the space environment, illustrating how large-scale clustering can reveal environment-specific adaptation. Similarly, subcluster analysis of 141,335 *Staphylococcus aureus* genomes recovered known clonal complexes, demonstrating that population structure appears to be preserved and recoverable at scale, but warrants further research beyond this single example.

We have several avenues for further improving performance and resolution. First, because *gemsparcl* uses a single k-mer size, and sketches capture a mixture of core-and accessory-genome signals, it prevents precise within-species resolution, although some is detected. A static ANI threshold cannot consolidate all genomes of highly diverse species into a single cluster. This is most extreme in *Helicobacter pylori*, where 4,833 genomes are distributed across 619 GCUs, reflecting the exceptional genomic diversity of this species. Similarly, *Salmonella enterica*, *Escherichia coli*, and *Campylobacter jejuni* each span multiple large GCUs despite representing a single species. Second, species that lack clear ANI boundaries, as described previously by Parks *et al*. as “fuzzy”, cannot be fully resolved by sequence similarity alone without anchoring to type-strain representatives. Addressing the first limitation is a priority for future development, as variable-length k-mers would enable large-scale within-species analysis and improve subspecies resolution. Querying new genomes against cluster representatives rather than all genomes could achieve near-linear time complexity, making incremental database updates computationally trivial as genome repositories continue to grow. Together, these developments would transform *gemsparcl* from a periodic clustering tool into a continuously maintained, queryable genomic index of bacterial diversity. Coupled to databases such as ENA, this could be used to flag inconsistencies in taxonomy and/or potential contamination.

As genomic databases continue to grow, tools that scale to their full complexity are not only convenient but a prerequisite for understanding the true breadth of microbial diversity.

## Supporting information

All supplementary figures are provided as a separate file

Supplementary Table 1: Parameter sweep for prefilter sketch size and k-mer length across combinations of k = 21, 61, 101, 301 and s = 3, 5, 8, 10, 20,

Singletons summary

Cluster summary

## Acknowledgements

Donovan H. Parks, Kate Ryan

## Author contributions

Conceptualisation: JvW, RDF, JAL, STH, Data curation: JvW, TAG, STH, Formal analysis: JvW, Funding acquisition: JvW, JAL, RDF, Investigation: JvW, Methodology: JvW, JAL, RDF, LJL, MJR, Project administration: RDF, JAL, Resources: JvW, TAG, Software: JvW, JAL, VRB, MJR, Supervision: RDF, JAL, STH, Validation: JvW, Visualization: JvW, Writing – original draft: JvW, JAL, LJL, Writing – review & editing: all

## Data and Code Availability

The entire genome clustering workflow: https://github.com/johannahelene/gemsparcl

The super fast sketching algorithm: https://github.com/bacpop/sketchlib.rust

Data: https://ftp.ebi.ac.uk/pub/databases/metagenomics/software/gemsparcl/

## Supplementary Materials

**Supplementary Figure 1:**
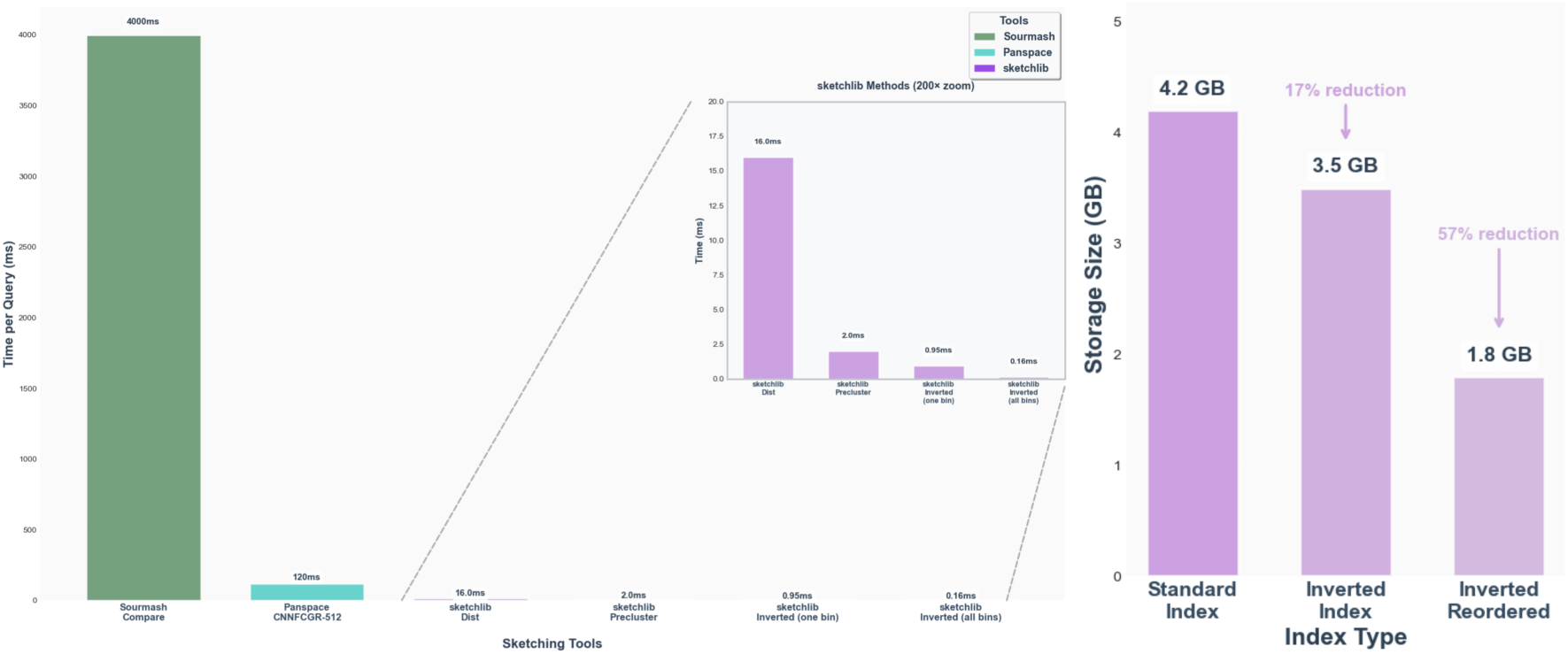
Query performance comparison across methods for the AllTheBacteria dataset (2.4M genomes, 32 threads). Wall-clock query time is shown for sourmash, Panspace (Cartes *et al*. 2025), and sketchlib (vanilla distances, prefiltered distances, and prefiltered distances with ordering) against the full AllTheBacteria dataset of 2,440,377 isolate genomes. Index sizes for standard, inverted, and reordered inverted sketch representations are shown for sketch size s=1000.

**Supplementary Figure 2:**
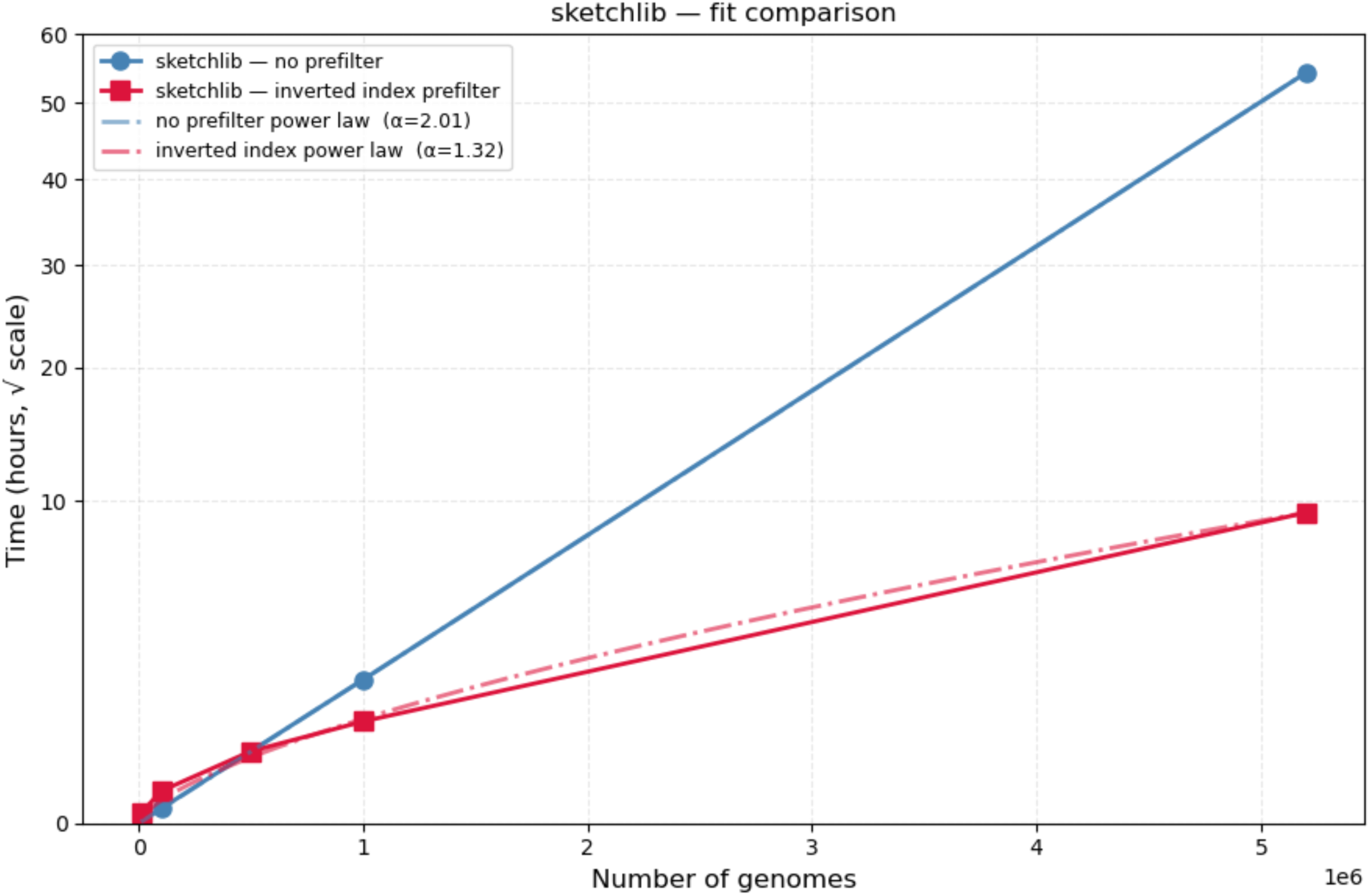
Runtime scaling of sketchlib with and without the inverted index. Wall-clock runtime is shown for standard sketchlib (blue) and the inverted index implementation (red) across increasing dataset sizes. Fitting a power law to each curve yields an exponent of α=2.01 for the standard implementation, confirming quadratic scaling, and α=1.32 for the inverted index implementation, demonstrating sub-quadratic scaling.

**Supplementary Figure 3:**
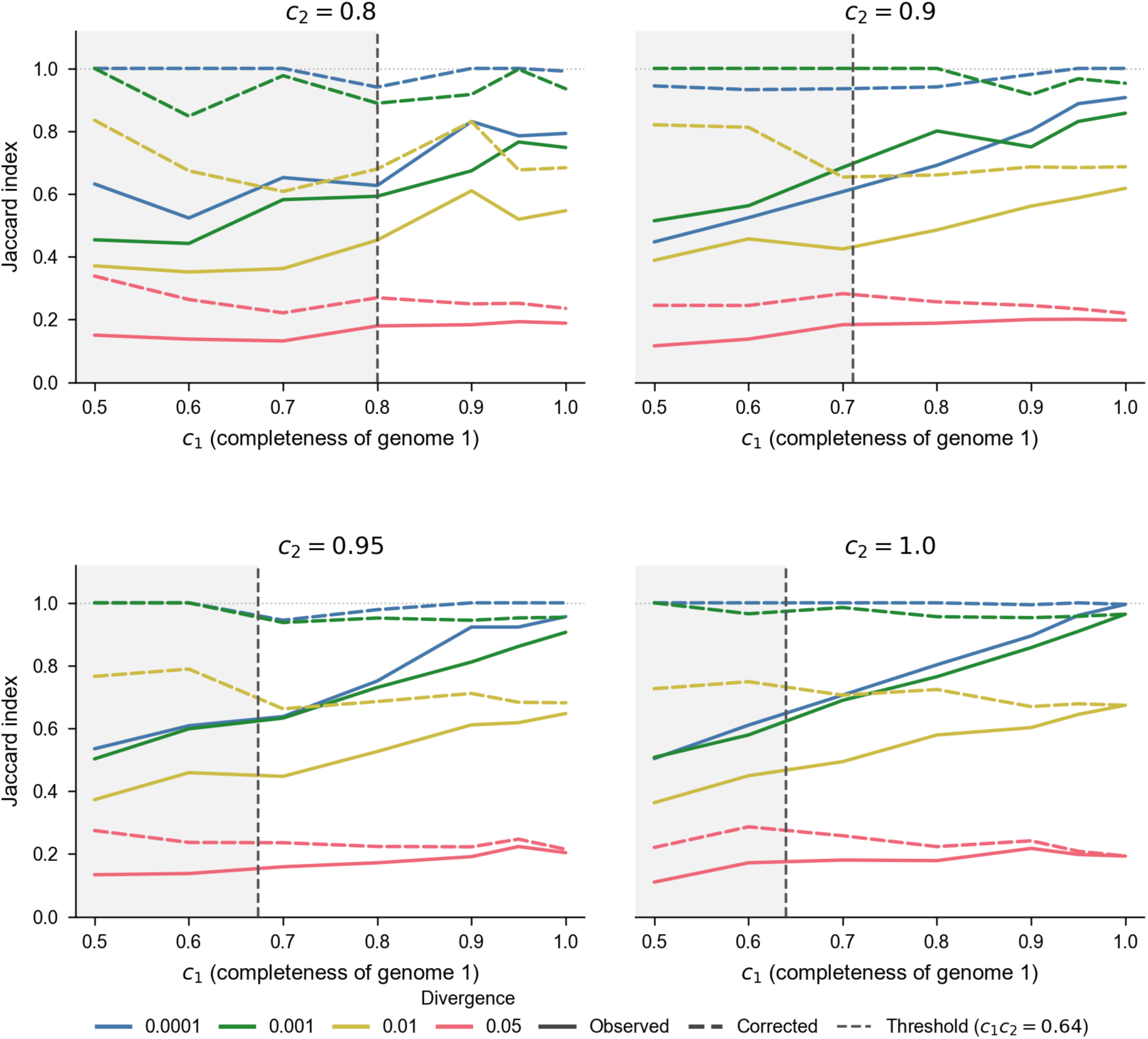
Genome incompleteness artificially deflates the Jaccard index and is corrected by completeness correction. Each panel shows a fixed completeness of genome 2 (c2 = 0.80, 0.90, 0.95, and 1.00), with the completeness of genome 1 (c1) varying along the x-axis. Within each panel, four lines represent genome pairs at increasing levels of divergence. For each divergence level, the observed Jaccard index (solid) is compared against the completeness-corrected Jaccard index (dashed), demonstrating that incompleteness systematically underestimates genomic similarity and that the correction restores accurate distance estimation across all tested completeness levels.

**Supplementary Figure 4:**
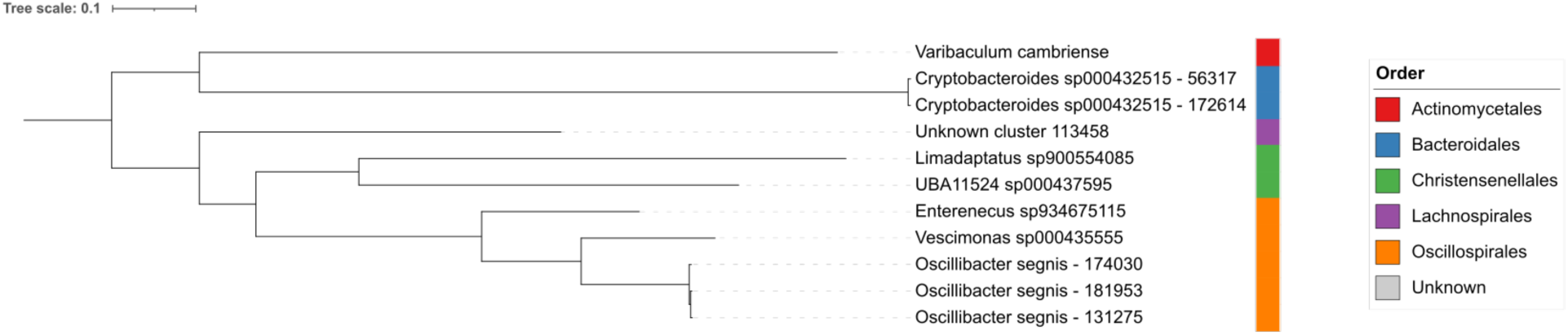
Maximum likelihood phylogeny of UBA11524 sp000437595 and its closest neighbouring species. The tree was inferred from GTDB-Tk marker gene alignments using IQ-TREE v3.0.1 (LG+G4 model, 1000 bootstrap replicates). Representative genomes were selected as the most complete genome from each neighbouring GCU. Branch lengths are in units of substitutions per site.

**Supplementary Figure 5:**
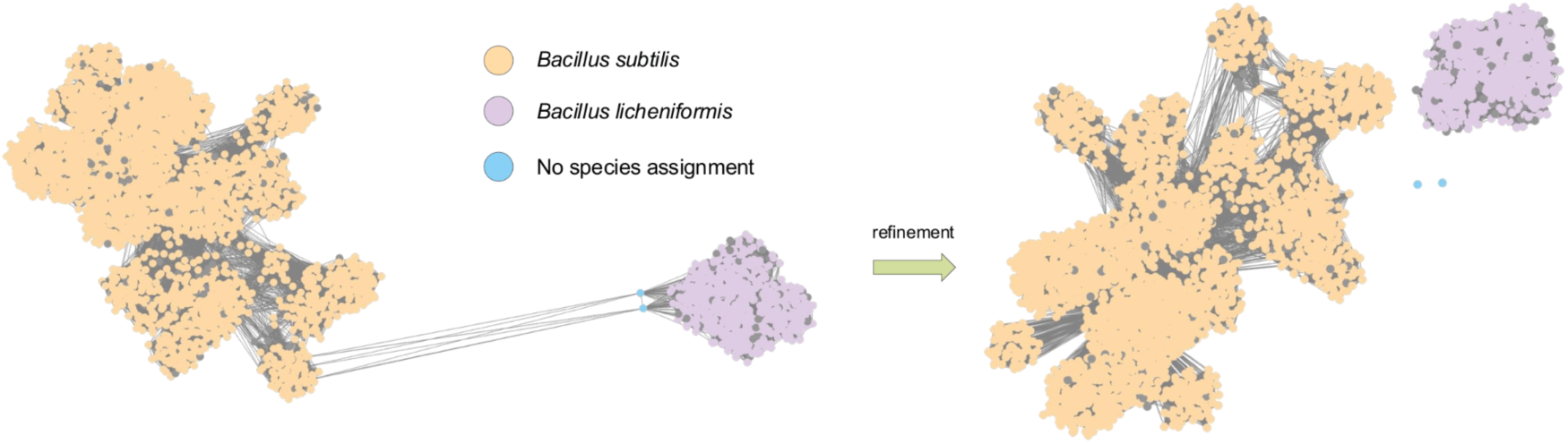
Example of refinement on GCU 155. Two similar species, *Bacillus subtilis* and *Bacillus licheniformis*, have been clustered together due to two bridge nodes (blue). These bridge nodes could not be taxonomically classified by GTDB-Tk, hence their blue colouring. The refinement algorithm identifies them, removes all their edges, and reintroduces them to the dataset as singletons.

**Supplementary Figure 6:**
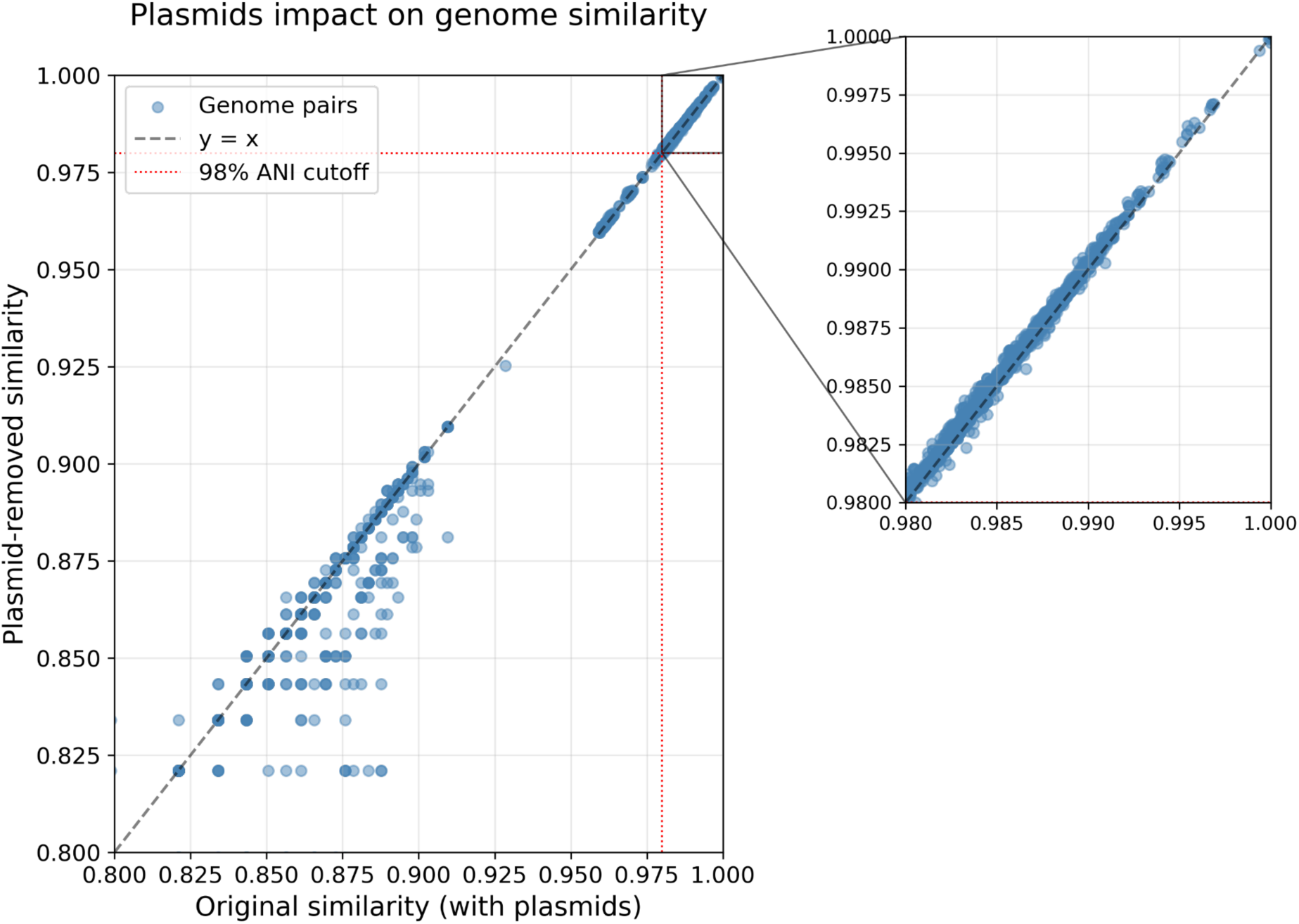
Plasmid content has minimal impact on genome similarity at clustering-relevant thresholds. Comparison of pairwise ANI values before and after plasmid removal for *Bacteroides uniformis* and *Phocaeicola vulgatus*. Genome pairs above 98% ANI (clustering threshold) align tightly with the perfect correlation line (dashed), demonstrating that plasmid sequences do not interfere with species-level clustering. Intra-species pairs at the top right match the correlated line, whereas for the inter-species correlation, the plasmids have a greater impact.

**Supplementary Table 1: Inverted index**

**Supplementary Table 2: Parameter sweep**

**Supplementary Table 3: Singletons**

**Supplementary Table 4: Cluster overview**

