## Supplementary material for "Rapid and consistent clustering of millions of genomes highlights the diversity of prokaryotic life": All supplementary figures are provided as a separate file

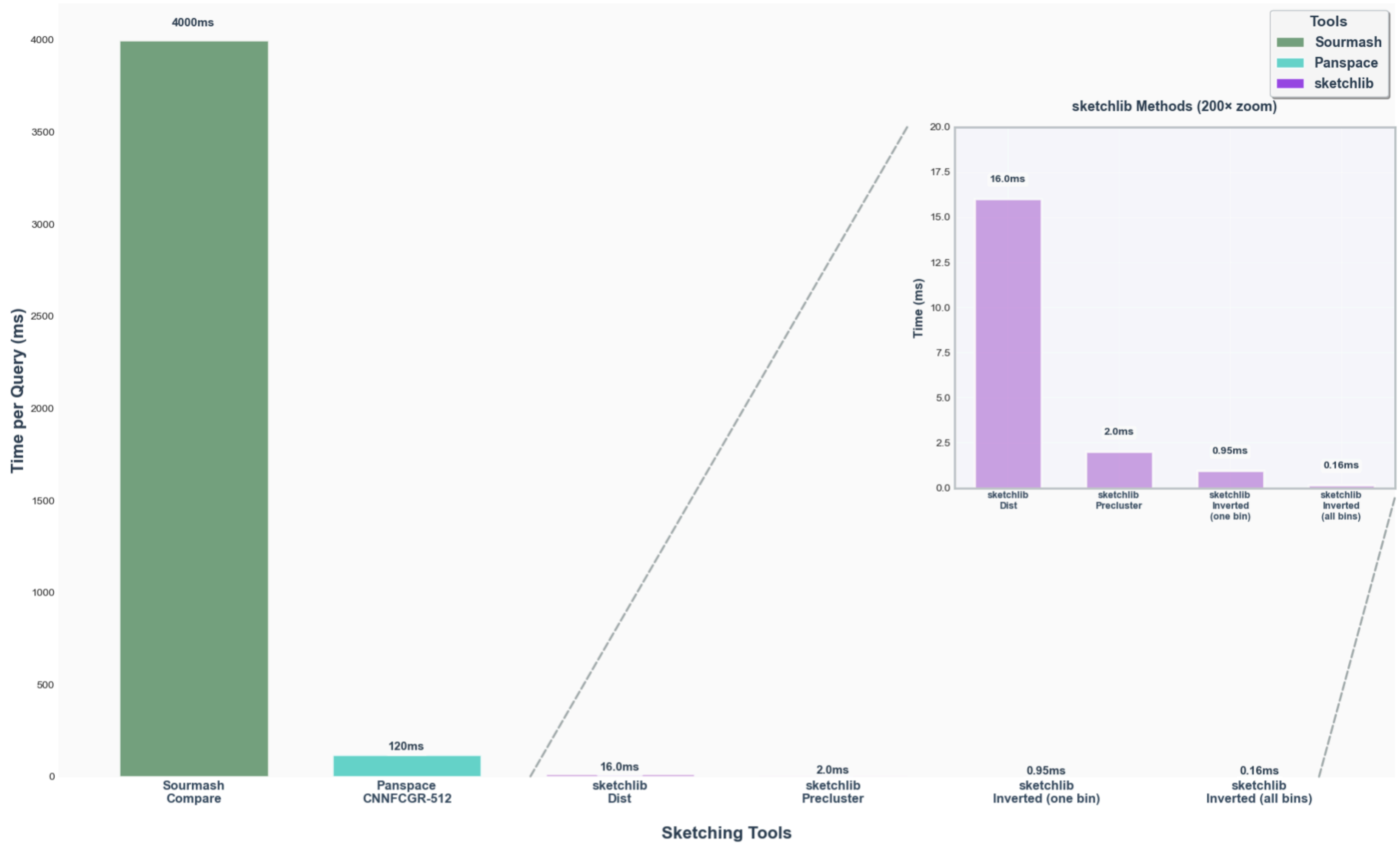
Time per Query (ms)
4000ms
120ms
16.0ms
2.0ms
0.95ms
0.16ms
Sourmash Compare
Panspace CNNFCGR-512
sketchlib Dist
sketchlib Precluster
sketchlib Inverted (one bin)
sketchlib Inverted (all bins)
Sketching Tools
Tools
Sourmash
Panspace
sketchlib
sketchlib Methods (200× zoom)
Time (ms)

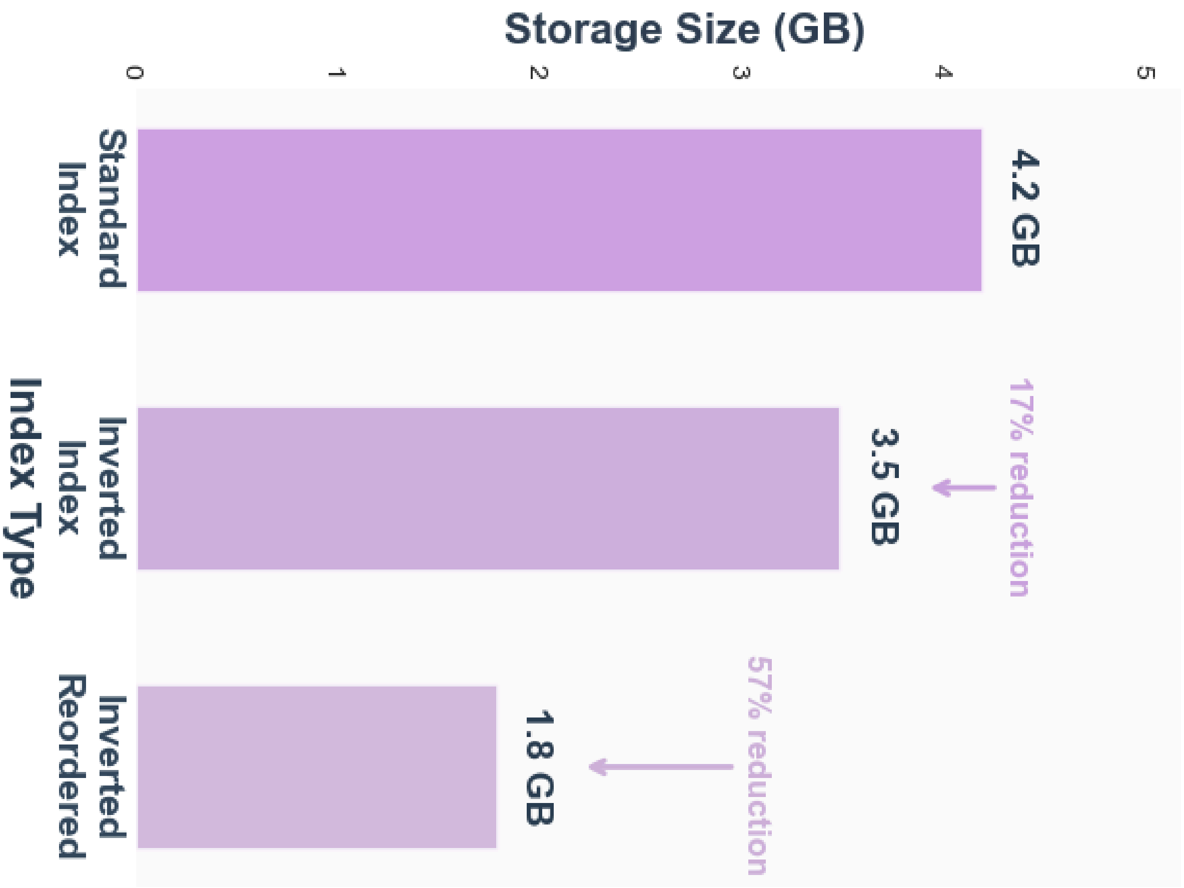


**Supplementary Figure 1: Query performance comparison across methods for the AllTheBacteria dataset (2.4M genomes, 32 threads).** Wall-clock query time is shown for sourmash, Panspace (Cartes *et al.* 2025), and sketchlib (vanilla distances, prefiltered distances, and prefiltered distances with ordering) against the full AllTheBacteria dataset of 2,440,377 isolate genomes. Index sizes for standard, inverted, and reordered inverted sketch representations are shown for sketch size s=1000.

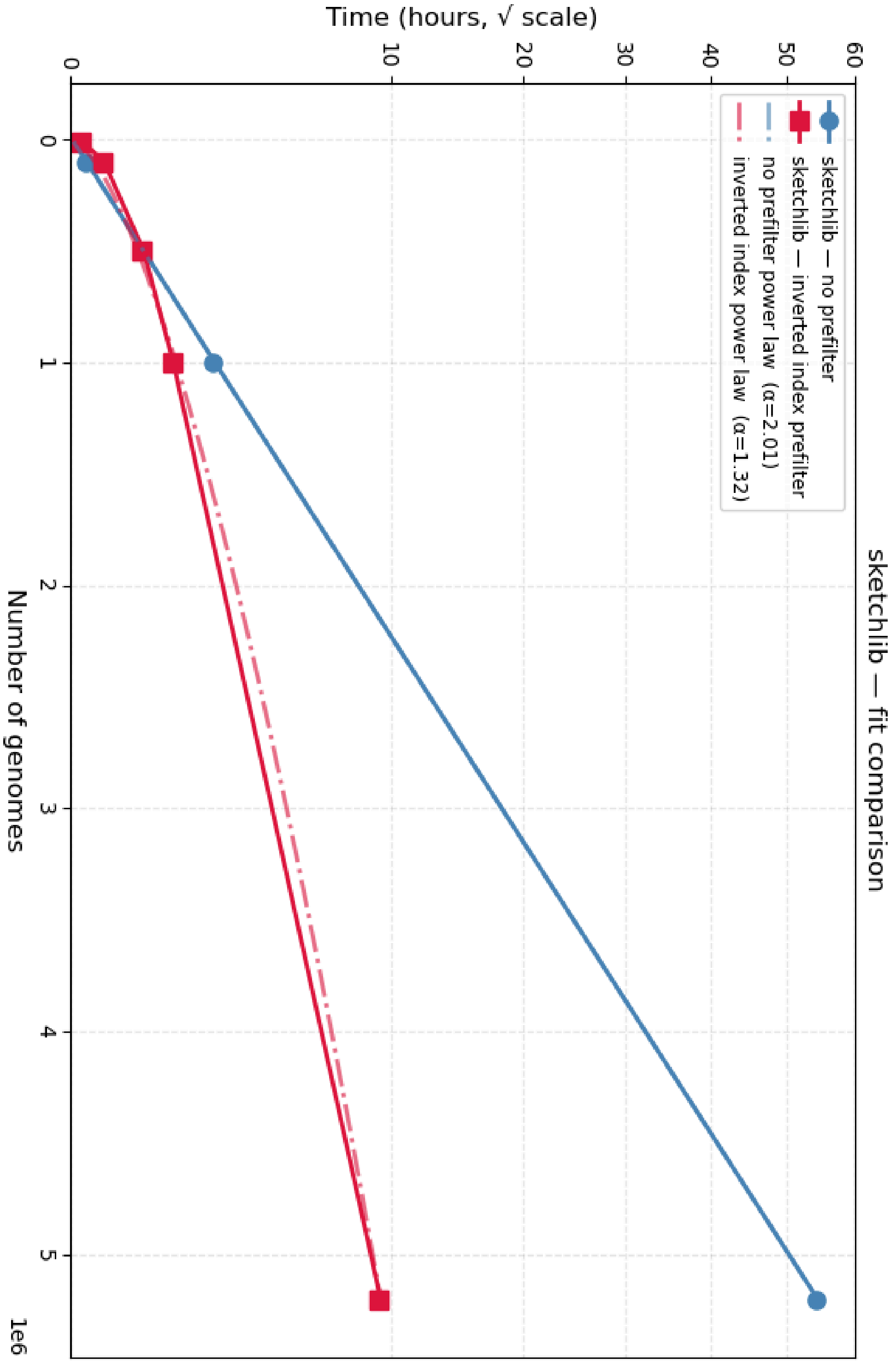


**Supplementary Figure 2: Runtime scaling of sketchlib with and without the inverted index.** Wall-clock runtime is shown for standard sketchlib (blue) and the inverted index implementation (red) across increasing dataset sizes. Fitting a power law to each curve yields an exponent of α=2.01 for the standard implementation, confirming quadratic scaling, and α=1.32 for the inverted index implementation, demonstrating sub-quadratic scaling.

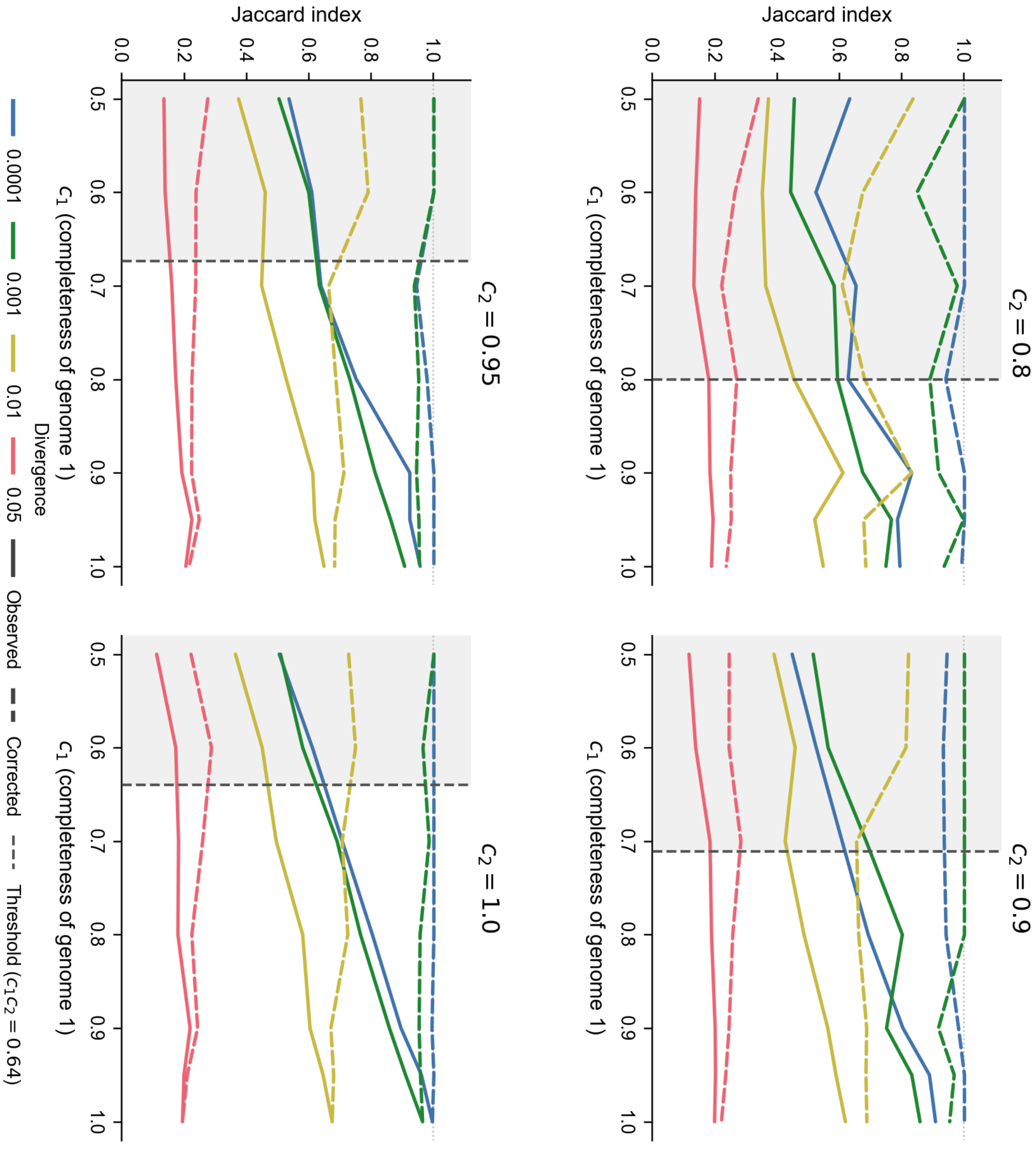


**Supplementary Figure 3: Genome incompleteness artificially deflates the Jaccard index and is corrected by completeness correction.** Each panel shows a fixed completeness of genome 2 (c2 = 0.80, 0.90, 0.95, and 1.00), with the completeness of genome 1 (c1) varying along the x-axis. Within each panel, four lines represent genome pairs at increasing levels of divergence. For each divergence level, the observed Jaccard index (solid) is compared against the completeness-corrected Jaccard index (dashed), demonstrating that incompleteness systematically underestimates genomic similarity and that the correction restores accurate distance estimation across all tested completeness levels.

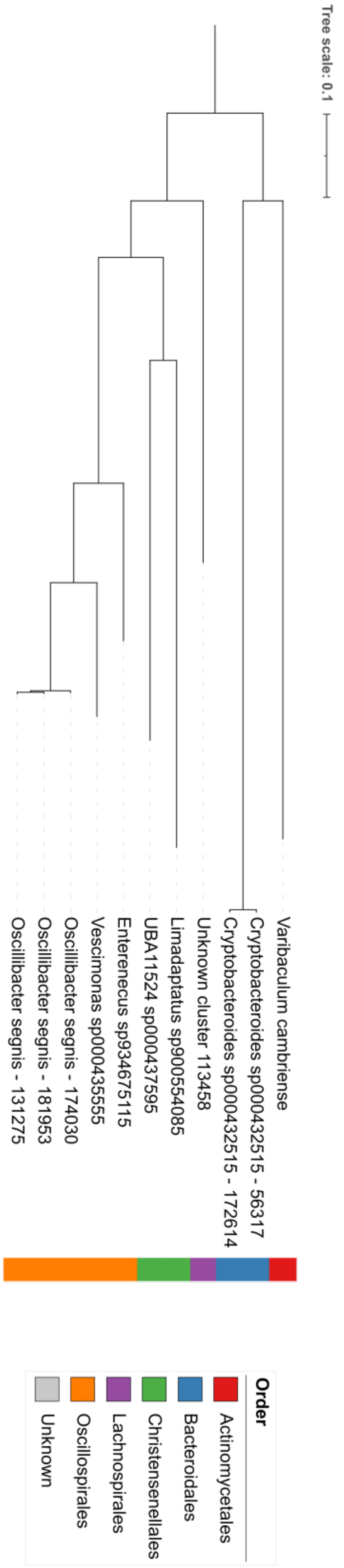


**Supplementary Figure 4: Maximum likelihood phylogeny of UBA11524 sp000437595 and its closest neighbouring species.** The tree was inferred from GTDB-Tk marker gene alignments using IQ-TREE v3.0.1 (LG+G4 model, 1000 bootstrap replicates). Representative genomes were selected as the most complete genome from each neighbouring GCU. Branch lengths are in units of substitutions per site.

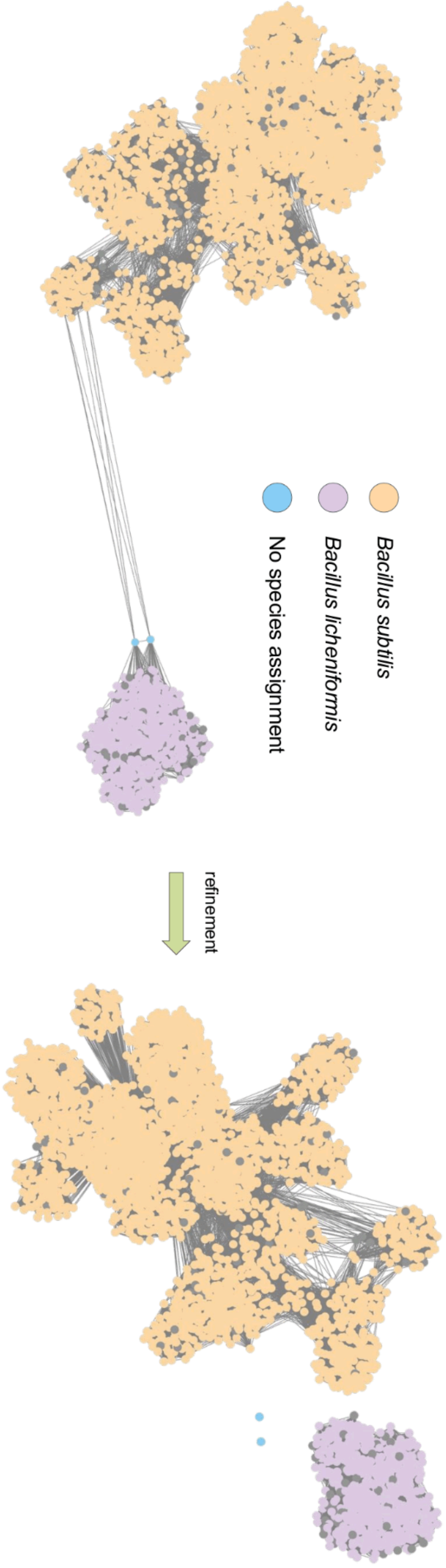


**Supplementary Figure 5:** Example of refinement on GCU 155. Two similar species, *Bacillus subtilis* and *Bacillus licheniformis*, have been clustered together due to two bridge nodes (blue). These bridge nodes could not be taxonomically classified by GTDB-Tk, hence their blue colouring. The refinement algorithm identifies them, removes all their edges, and reintroduces them to the dataset as singletons.

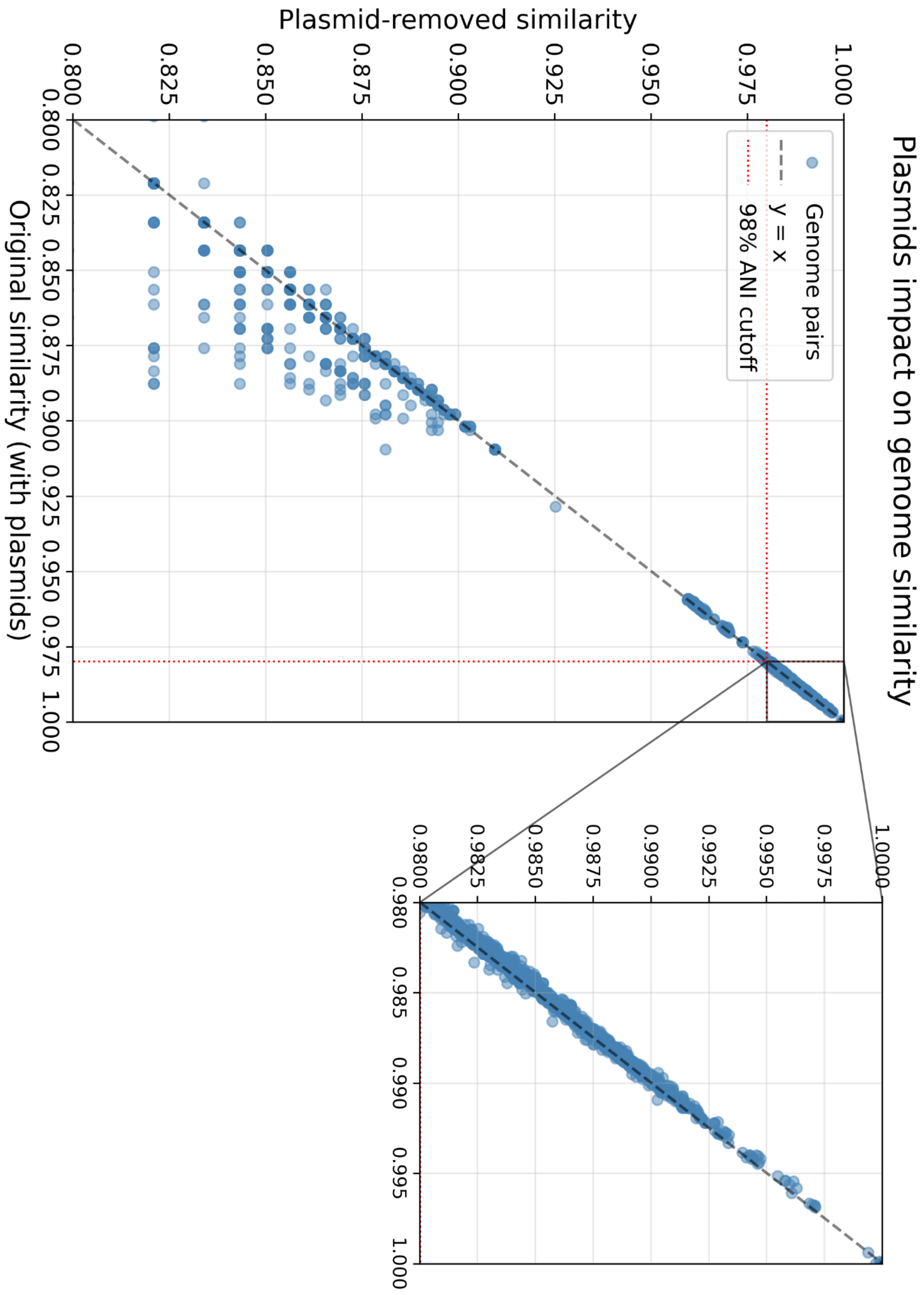


**Supplementary Figure 6: Plasmid content has minimal impact on genome similarity at clustering-relevant thresholds.** Comparison of pairwise ANI values before and after plasmid removal for *Bacteroides uniformis* and *Phocaeicola vulgatus*. Genome pairs above 98% ANI (clustering threshold) align tightly with the perfect correlation line (dashed), demonstrating that plasmid sequences do not interfere with species-level clustering. Intra-species pairs at the top right match the correlated line, whereas for the inter-species correlation, the plasmids have a greater impact.
